# The oncogenic CCDC6-RET fusion product is a dual ATP and ADP-dependent kinase that functions via *cis*-phosphorylation

**DOI:** 10.1101/2025.01.14.632894

**Authors:** Ana Martín-Hurtado, Julia Contreras, Jana Sánchez-Wandelmer, Eduardo Zarzuela, Inés G. Muñoz, Javier Muñoz, Marta Isasa, Iván Plaza-Menacho

**Affiliations:** Protein Phosphorylation and Cancer Group, Structural Biology Programme; Proteomics Unit; Protein Crystallography Unit Spanish National Cancer Research Center (CNIO) C/Melchor Fernández Almagro num. 3, 28029 Madrid, Spain (ext. 3030)

## Abstract

Gene fusions products involving protein kinases are known drivers in human cancers and actionable targets for personalized therapy, yet the structural and molecular determinants that control their function are largely unexplored. Here we show that a CCDC6-RET fusion product, a driver and therapeutic target in lung and thyroid cancers, is a highly active dimeric kinase in solution. Time-resolved mass spectrometry analysis together with a robust biochemical and biophysical characterization reveal that CCDC6-RET functions as a dual ATP- and ADP-dependent kinase able to bind both nucleotides and uses them as phosphoryl donors. We also identify a crosstalk between the c-terminal and the activation segments, uncovering a mutually exclusive dependency by the former on activation loop phospho-sites controlling both the processing and the catalytic activity of the fusion protein. Furthermore, we generated a 3D-assembly of a CCDC6-RET homodimer combining electron microscopy (EM) single particle, small-angle X-ray scattering (SAXS) and in silico structural analyses. Our structural model together with cross-linking mass spectrometry data demonstrated that CCDC6-RET forms a face-to-face *trans*-inhibited dimer in the apo state characterized by intermolecular-crosslinked activation segments. Upon nucleotide binding and catalytic domains reorientation, fast activation loop phosphorylation is driven by a mechanism *in cis*. Our work uncover for the first time the molecular and structural determinants that controls CCDC6-RET function and provides a solid framework to study the structure and function of other RET fusion products.

## Introduction

Following the paradigm of the Philadelphia chromosome and the BCR-ABL1 fusion in chronic myelogenous leukemia (CML) (Nowell, 1960) (de Klein et al., 1982), gene fusions products involving protein kinases are known drivers in human cancers and actionable targets for personalized therapy (Santoro et al., 2020). Yet, the structural and molecular determinants that control their function and (oncogenic) mechanism of action are in most of the cases paradoxically elusive. In 1985, a novel proto-oncogene named Rearranged during Transfection (RET) was discovered in NIH3T3 cells transfected with DNA isolated from human lymphoma cells, which resulted in malignant transformation (Takahashi et al., 1985). The characterization of the isolated *RET* cDNA clones revealed a carboxy-terminal domain with high homology with members of the tyrosine kinase gene family preceded by a hydrophobic sequence characteristic of a transmembrane domain, suggesting that the *RET* oncogene encoded a cell-surface receptor (Takahashi and Cooper, 1987). The characterization of the human (Takahashi, 1988) and mouse (Iwamoto et al., 1993; Pachnis et al., 1993) proto-oncogene orthologs revealed the amino acid sequence (primary structure) of the RET protein. Shortly after, a RET fusion product was discovered in papillary thyroid carcinoma (PTC) named RET/PTC1 and its N-terminal partner protein identified as H4(D10S170) (Fusco et al., 1987; Grieco et al., 1990). The RET/PTC-1 oncogene was a fusion product containing the n-terminus of the H4 gene, currently known as CCDC6 (Coiled-Coil Domain Containing protein 6, aa 1-101) fused to the tyrosine kinase domain of c-RET (Grieco et al., 1990). CCDC6-RET fusions are detected in up to 30% of papillary thyroid carcinomas (Grieco et al., 1990; Jhiang et al., 1992). Although prevalence of CCDC6-RET fusions in PTC shows significant geographic variation (Nikiforov, 2002), there is a strong correlation with radiation exposure. In 1986, after the Chernobyl nuclear power disaster, a sharp increase in the incidence of PTC was observed in children and adolescents of contaminated regions of Belarus, Russia, and Ukraine with near 60% of post-Chernobyl PTCs children patients harboring a RET fusion (Ricarte-Filho et al., 2013). Up to date, more than twenty RET fusions products have been described in PTC (Santoro et al., 2020); the most common RET fusions products are CCDC6-RET (RET/PTC1) and NCOA4-RET (RET/PTC3), accounting for about 90% of RET fusion-positive cases in thyroid cancer (Nikiforov, 2002). In all these cases, the DNA-rearrangements generate the fusion of the tyrosine kinase domain of RET to the 5’ portion of different genes. Breakpoints in the RET gene are consistently located at intron 11, so the fusion starts from exon 12 to the stop codon, resulting in a protein chimera containing the tyrosine kinase domain without the juxtamembrane segment nor extracellular domains (Nikiforov, 2002). A common molecular hallmark in PTC is the constitutive activation of the MAPK (Mitogen-Activated Protein Kinase) cascade. Indeed, besides RET and other RTKs, common PTC-driver are gain-of-function mutations of RAS and BRAF kinase, respectively. The mutually exclusive nature of these lesions supports the notion they act along a common signaling axis driving the oncogenic phenotype in PTC (Fagin and Wells, 2016). RET fusions are also found in 1-2% of lung adenocarcinoma (LADC) (Kohno et al., 2012; Lipson et al., 2012; Wang et al., 2012). The most common RET fusion products in NSCLC are KIF5B-RET, NCOA4-RET, and CCDC6-RET respectively. KIF5B (Kinesin-1 heavy chain) is by far the most common RET fusion partner in NSCLC, being detected in up to 70%–90% of the cases (Kohno et al., 2012). In NSCLC, RET fusions are mutually exclusive with other driver mutations, such as ALK or ROS1 rearrangements or EGFR or KRAS mutations, suggesting again common signalling mechanisms driving the oncogenic phenotype, and the definition of a potential molecular subtype (Klempner et al., 2015).

Clinically, there is strong interest in the development of next generations RET tyrosine kinase inhibitors for the development of personalized therapies in NSCLC and PTC (Shehata et al., 2023). Crucially RET fusions (CCDC6-RET, NCOA4-RET, and the newly described CDC123-RET) have been shown to mediate acquired resistance mechanism in lung adenocarcinomas (LADC) to EGFR or ALK tyrosine kinase inhibitors (de Klein et al., 1982; McCoach et al., 2018; Piotrowska et al., 2018). The understanding of the structural, molecular and dynamic determinants that control the catalytic function and signalling properties of these oncogenic RET fusion products will uncover druggable vulnerabilities that can be therapeutically exploited for the design and development of next generation inhibitors for targeted and personalized therapy that are able to overcome acquired resistance by secondary mutations (Lin et al., 2020; Shehata et al., 2023; Solomon et al., 2020; Tan and Solomon, 2020; Terzyan et al., 2019).

In this study we produced in Sf9 insect cells using a baculovirus system a recombinant CCDC6-RET gene fusion product. We performed a detailed functional and structural characterization of the CCDC6-RET protein by state-of-the-art biochemistry, mass spectrometry and structural biology tools that allowed us to generate a 3D-structural model of CCDC6-RET homodimer for single particle analyses and mechanistic validation. Our findings uncover for the first time the structural and molecular determinants that control a RET oncogenic fusion’s mechanism of autoactivation, revealing unexpected and striking features not previously envisioned with important implication for drug-discovery. Our data provides at the same time, a solid framework for the structural and functional study of other RET fusion products.

## Materials and methods

### Plasmids

A modified pBac-PAK3 plasmid (Invitrogen) containing an N-terminal 3 x His and a GST (Glutathione S-transferase) tags followed by a Human Rhinovirus (HRV)-3C protease cleavage site was used to express a codon optimized CCDC6-RET fusion product composed by fragments of residues 1-101 of human CCDC6 and 713-1072 of human RET (Knowles et al., 2006; Plaza-Menacho et al., 2016; Plaza-Menacho et al., 2014a). A construct lacking the c-terminal segment of RET (RET aa 713-1013, i.e. ΔCT) was also generated. These plasmids were used to create different point mutants by site-directed mutagenesis.

### Site-directed mutagenesis

Site-directed mutagenesis was performed by using Q5-site directed mutagenesis kit (New England Biolabs) following manufacturer instructions and the indicated non-overlapping primers:

RET_Y900F- forward 5’- CCGTGACGTGTTCGAGGAGGACT -3’

RET_Y900F- reverse 5’- CTCAGTCCGAAGTCGGAG -3’

RET_Y905F- forward 5’- GGAGGACTCCTTCGTCAAGCGTT -3’

RET_Y905F- reverse 5’- TCGTACACGTCACGGCTC -3’

RET_Y1015F- forward 5’- CGTCGTGACTTCCTGGACCTG -3’

RET_Y1015F- reverse 5’- CTTGACCATCATCTTCTCCAG -3’

RET_Y1062F- forward 5’- AACAAGCTCTTCGGTCGTATCTC -3’

RET_Y1062F- reverse 5’- CTCGATCCAGGTGGAGGG -3’

RET_S1034A_forward 5’- CGACGGACTGGCCGAGGAGGAGA -3’

RET_S1034A_reverse 5’- TCGTAGATCAGGGAGTCGGAGG -3’

RET_S98A_forward 5’-GCGCAAGGCTGCGGTGACTATCG -3’

RET_S98A_reverse 5’- AGGTCGCGGTTTTCC -3

CCDC6_S52A forward 5’- AATCGTGATCGCCCCATTCCGTC -3’

CCDC6_S52A reverse 5’- CCACCAGATTTTCCACCG -3’

CCDC6_C84A_forward 5’- CAAGCTGAAGGCCAAGGCTCTGC -3’

CCDC6_C84A_reverse 5’- TAGGTTTCCAGCTCGATC -3’

CCDC6_Y80F_forward 5’- GCTGGAAACCTTCAAGCTGAAGT -3’

CCDC6_Y80F_reverse 5’- TCGATCTTCAGCACCTTG -3’

Alternatively, other mutations were generated by using overlapping primers:

RET_S909A forward 5’- CGTCAAGCGTGCCCAGGGTCGTA -3

RET_S909A reverse 5’- TACGACCCTGGGCACGCTTGACG -3’

CCDC6_S46A forward 5’- CGGTGGAAAAGCTGGTGGAATCGTGATCTC -3’

CCDC6_S46A reverse 5’- GAGATCACGATTCCACCAGCTTTTCCACCG -3’

### Heterologous protein expression in insect cells

We used the *flash*BAC^TM^ system (Oxford Expression Technologies) for recombinant protein expression using a baculovirus system (Knowles et al., 2006; Plaza-Menacho et al., 2016; Plaza-Menacho et al., 2014a) following the manufacturer recommendations. Briefly, Sf9 insect cells were seeded in a 6 wells/plate at a density of 0.3-0.5 x 10^6^ cells/mL in serum free Sf900-II media (Gibc*o*) supplemented with 200 μL of Gentamycin (50 mg/mL) per liter. Next, the transfection reaction was prepared in a sterile Eppendorf tube containing antibiotic-free media Sf900-II (up-to 1 mL) and the transfection mixture composed by: 2.5 μL of *flash*Bac^TM^ linear viral DNA (20 ng/ μL), 1 μL (500 ng) of transfer plasmid (pBacPAK3) and 5 μL of transfection reagent (Fugene) following manufacturer instructions. After 5-6 days, the supernatant of the transfected cells was collected to have the recombinant baculovirus stock (P1). The P1 baculovirus was inoculated (1:25) in a suspension culture of 25 mL of complete Sf900-II media at a density of 1.5-2 x 10^6^ cells/ml and incubated shaking at 90-95 rpm for 3-4 days. After this time, the supernatant was recovered obtaining the P2 baculovirus. After testing the expression of the protein, the P2 baculovirus can be used to inoculate another 100 mL suspension culture of Sf9 cells as indicated before at a 1:100 dilution. After 3-4 days (72-96 hours), the supernatant became the P3 baculovirus. Both, the P2 and P3 baculovirus were used as baculovirus stocks for protein production.

### Protein purification

Protein purification was achieved by in tandem IMAC (Ni^+2^) and Glutathione-conjugated gravity flow chromatography and in-gel 3C-protease digestion followed by size exclusion chromatography if required. This optimized protocol resulted in highly pure and monodisperse recombinant fusion product. Inoculated insect cell cultures (72-96 hours) were harvested by centrifugation at 3000 rpms (10 min, 4°C). Pellets were stored at −80 °C or used directly for protein purification. Pellets were resuspended in lysis buffer (50 mM Tris pH 7.5, 500 mM NaCl, 10 mM Benzamidine, 0.2 mM AEBSF, 2 mM TCEP). After sonication on ice (20 cycles 9 seconds on/ 3 seconds off and 37% amplitude), crude lysates were centrifugated at 20000 rpms at 4 °C for 45 min. Clarified lysate underwent a final sonication step of 10 sec followed by filtration with a 45 μm filter (Jet Biofilter) with a syringe. Crude lysate, clarified lysate (supernatant) and unsoluble material (pellet) samples were mixed with 5X Laemmli sample buffer (*ThermoFisher*) and run on SDS-Polyacrylamide gel electrophoresis (SDS-PAGE). First, the clarified lysate was incubated with a previously equilibrated Nickel (Ni^2+^) chelate resin. After two hours incubation on rotation in a cold room, the flow through (FT) was separated and beads were washed with Buffer A (20 mM Tris pH 7.5, 500 mM NaCl, 2 mM TCEP) with additional 20 mM Imidazole. Protein elution was done with Buffer B (20 mM Tris pH 7.5, 500 mM NaCl, 400 mM Imidazole, 2 mM TCEP) using a filter spin column (*SC 1000-1KT, Lot SLCD4489, SIGMA-ALDRICH*) by centrifugation at 8.000 rpm during 2 min. Fractions were checked by SDS-PAGE stained with Coomassie previously to the second purification step. Positive fractions eluted from the His-affinity step were incubated with Glutathione conjugated Sepharose beads (*Glutathione Sepharose^®^ 4 Fast Flow, Cytiva*) for several hours at 4 °C. Next, the GST-tagged protein was separated by centrifugation (900 rpm) and washed 4-5 times with 10 mL of GST-Buffer (20 mM Tris pH 7.5, 500 mM NaCl, 2mM DTT) prior to in gel digestion (4 hours) with an GST-3C protease (200 μg/L culture) and elution (2-3) using a filter spin column (8.000 rpm, 2 min). Eluted fractions were checked by SDS-PAGE gel stained with Coomassie. Protein concentration and purity were measured by absorbance using a Nanodrop. When required a size exclusion chromatography (SEC) step using AKTA pure protein purification system (*GE Healthcare*) was performed. The sample was concentrated up to 2 mL with a Millipore concentrator with a 30 kDa cut-off by centrifugation (Eppendorf Centrifuge 5810 R) at 2.000 rpms at 4 °C. The concentrated sample was injected into a Superdex 200 16/60 or 10/300 columns (*GE Healthcare*). The input of the column together with the collected fractions were analyzed by SDS-PAGE followed by Coomassie staining, and the protein concentration was measured by absorbance.

### SDS-PAGE, western blotting (WB) and antibodies

Proteins were electrophoretically separated on 12% SDS-PAGE gels run in the Mini-PROTEAN system with electrophoresis buffer (25 mM Tris pH 8.3, 192 mM glycine, 0.1% SDS) by using a BIO RAD PowerPac Basic at a constant voltage of 150-200 mV for 30-45min. Pre-stained dual colour markers (*Precision Plus Protein Standards* (10-250 kDa)*, BIO-RAD and Protein Ladder PS10 Plus (11-180 kDa), GeneOn*) were used as molecular weight standards. For protein staining, gels were gently shacked in Coomassie staining solution (10% acetic acid, 40% absolute ethanol and 50% MilliQ water with 1g/L Brilliant blue 250 R) for 30-60 min. Then the gels were distained for 1-2 hours with a distaining solution (10% acetic acid, 50% absolute ethanol, 40% MilliQ water). Proteins were transferred from SDS-PAGE gels onto nitrocellulose 0.2 μm (*Amersham*) or PVDF 0.45 μm (Millipore) membranes using the Mini-PROTEAN system (*BIO RAD*). Before transference, PVDF membranes were activated first in ethanol 100% for 10 min. Membrane were wet-transferred in transfer buffer (25 mM Tris pH 8.8, Glycine 190 mM and ethanol 10%) on ice at constant 100-200 V for 60 min. Next, transferred membranes were blocked in TBS (10 mM Tris pH 8, 150 mM NaCl) supplemented with 5% (w/v) skimmed powder milk and gently shacked for 30 min. After blocking step, membranes were washed in TBS-T (10 mM Tris pH 8, 150 mM NaCl Tween-20 0.1% v/v) for 3-4 times. Then they were incubated with constant agitation (*Duomax 1030, Heidolph*) O/N at 4°C with the following primary antibodies diluted in TBS-T supplemented with 5% BSA (w/v): phospho-Tyr (pTyr-100, CST #9411), RET (D3D8R, CST #14698), phospho-Ret Tyr 905 (CST #3221), phospho-Ret Tyr 1015 (Abcam # 74154) and phospho-Ret Tyr 1062 (Abcam # 51103) at 1:5.000 or 1:10.000 dilution. After overnight incubation with the primary antibody, membranes were washed with TBS-T for 3-4 times and incubated with the secondary antibody for 60 min with gently shaking protected from light. Secondary antibodies used were: anti-rabbit IgG (H+L) conjugated at 800 nm (*DyLight 800 Conjugate #5151, CST*) and anti-mouse IgG (H+L) conjugated at 680 nm (*DyLight 680 Conjugate #5470, CST*). These secondary antibodies were diluted 1:20.000 in TBS-T with 5% (w/v) skimmed powder milk. After 60 min of secondary antibody incubation, membranes were washed with 20mL of TBS-T for 3-4 times. Finally, proteins were detected in the membranes with the Odyssey CLx scanner (*LI-COR Biosciences*) and the images were analyzed with the *Image Studio lite* software.

### Differential Scanning Fluorometry (DSF)

To determine the thermal stability of the recombinants proteins we apply a direct method based on intrinsic fluorescence (IF) detection using a Tycho NT.6 instrument (*NanoTemper Technologies*). Briefly, a protein sample (1-2 μM, 12μL) was loaded into a glass capillary tube and heated from 35 °C to 95 °C at a rate of 30 °C /min. The inflection temperatures (T_i_) were calculated by the instrument based on the 350/330nm fluorescence ratio. We measure the increment in thermal stability in absence or presence of different ligand and inhibitors, as indicated in the text.

### Mass Photometry

To determine the molecular mass distributions (40 kDa to 5 MDa) of single molecules we applied light scattering to detect individual, unlabeled molecules in dilute solutions using a MPone instrument (*Refeyn*). All the events were recorded with *Refeyn Acquire MP* software and analyzed with *Refeyn Discover MP* software to obtain mass distribution of protein samples.

### In vitro phosphorylation assays

For autophosphorylation experiments, recombinant protein 1-2 μM (final concentration) was used in 20 mM Tris pH 7.5, 500 mM NaCl, 2mM DTT and 2 mM MgCl_2_. We performed a time-points reaction triggered by the addition of ATP (1 mM): unstimulated, 1, 5, 15, 30 min. To stop the phosphorylation reaction samples were mixed with 5X Laemmli sample buffer and denaturalized at 95°C during 2-5 min in a thermoblock (*AccuBlock, Labnet*). In the case of phosphorylation experiments using intact susbtrate surrogates (catalytic dead) a 1:3 molar excess of the susbtrate to the enzyme was employed in the reaction.

### Enzyme kinetic assays

Rates of phospho-tyrosine activity by recombinant CCDC6-RET and its variants were determined by a continuous ATP-coupled pyruvate kinase essay based on the consumption of NADH in the absence and presence of different peptides susbtrates at increasing concentrations of ATP (Cuesta-Hernandez et al., 2023; Plaza-Menacho et al., 2016; Plaza-Menacho et al., 2014a). The enzyme-substrate solution (1 mM Tris-Cl pH 7.5, 1 mM MgSO_4_, 400 mM phosphoenol pyruvate (PEP), 100 mM NADH, 2450 U/mL Pyruvate kinase, 2.26 mg/mL Lactate dehydrogenase) was prepared at different concentrations of ATP (mM): 0, 0.08, 0.16, 0.32, 0.65, 1.25, 2.5 and 5. Unless otherwise stated, the final protein concentration used in these assays was 0.2-1 μM, and the concentration of the peptides used EAIYAAPFAKKK and the RET A-loop DVYEEDSYV were 1.35 and 0.55 mM, respectively. The experiments were performed in a 384 well plate (*Greiner bio-one*) and the NADH consumption was read at 340 nm wavelength during 120 cycles of 30 sec each by using a microplate reader (*CLARIOstar Plus, BMG LABTECH).* NADH consumption curves were obtained and slopes were calculated by using de *CLARIOstar Data Analysis* software. Finally, by using *Prism* software, catalytic rates and kinetic constants were obtained by fitting the data to the Michaelis-Menten equation.

### Mass Spectrometry

For liquid chromatography coupled to tandem mass spectrometry (LC-MS/MS) samples were reduced and alkylated in 15 mM TCEP, 30 mM CAA, 30 min at RT in the dark in the presence of urea 4 M and digested with trypsin (Promega) in urea 1 M in 50 mM Tris pH 8 overnight at 37 °C at an estimated protein:enzyme ratio of 1:100. Digestion was quenched by 0.1% TFA and resulting peptides were desalted using C18 stage-tips. LC-MS/MS was performed by coupling an UltiMate 3000 RSLCnano LC system to a Q Exactive Plus mass spectrometer (*Thermo Fisher Scientific*). Digested peptides were loaded into a trap column (Acclaim™ PepMap™ 100 C18 LC Columns 5 µm, 20 mm length) for 3 min at a flow rate of 10 µL/min in 0.1% formic acid. Then, peptides were transferred to an EASY-Spray PepMap RSLC C18 column (*Thermo*) (2 µm, 75 µm x 50 cm) operated at 45 °C and separated using a 60 min effective gradient (buffer A: 0.1% FA; buffer B: 100% ACN, 0.1% FA) at a flow rate of 250 nL/min. The gradient used was, from 4% to 6% B in 2 min, from 6% to 33% B in 58 min, plus 10 additional min at 98% B. Peptides were sprayed at 1.5 kV into the mass spectrometer via the EASY-Spray source. The capillary temperature was set to 300 °C. The mass spectrometer was operated in a data-dependent mode, with an automatic switch between MS and MS/MS scans using a top 15 method (intensity threshold ≥ 4.5 x 10^4^, dynamic exclusion of 5 or 10 sec and excluding charges unassigned, +1 and > +6). MS spectra were acquired from 350 to 1500 m/z with a resolution of 70,000 FWHM (200 m/z). Ion peptides were isolated using a 2.0 Th window and fragmented using higher-energy collisional dissociation (HCD) with a normalized collision energy of 27. MS/MS spectra resolution was set to 35,000 (200 m/z). The ion target values were 3 x 10^6^ for MS (maximum IT of 25 ms) and 1×10^5^ for MS/MS (maximum IT of 110 msec).

Raw files were processed with Maxquant (v 1.6 and higher) using the standard settings against the corresponding sequences of the expressed recombinant proteins. Carbamidomethylation of cysteines was set as a fixed modification whereas oxidation of methionines, acetylation and phosphorylation of serines, threonines and tyrosines were set as variable modifications. Minimal peptide length was set to 7 amino acids and a maximum of two tryptic missed-cleavages were allowed. Results were filtered at 0.01 FDR (peptide and protein level). Raw data were imported into Skyline. Label free quantification of identified phosphopeptides was performed using the extracted ion chromatogram of the isotopic distribution. Only peaks without interference were used for quantification. Phosphopeptides intensities were normalized by the intensity of non-modified peptides from the target protein.

### Crosslinking mass spectrometry

Both, phosphorylated (1 mM ATP, 2 mM MgCl_2_, 30 min) and un-phosphorylated CCDC6-RET (4-6 µM) were crosslinked with 200-fold molar excess of disuccinimidyl dibutyric urea (DSBU, Thermo scientific) in 20 mM HEPES pH 7.5, 500 mM NaCl, 1mM TCEP. All the crosslinking reactions were performed at RT for 50 min and then quenched with 20 nM NH_4_HCO_3_. The cross-linked samples were vacuum-dried and resuspended in 4% SDS in 50 mM Tris/HCl pH 8.0. Reduction and alkylation were performed with 15 mM TCEP and 25 mM chloroacetamide 30 min at 45 °C, respectively. Samples were then trypsinized (16 hours, 37 °C) at an enzyme-to-substrate ratio of 1:50 following an automated SP3 protocol (Hughes et al., 2019). Digested samples were adjusted to 20 mM NH_4_OH and fractionated in four fractions by micro reverse phase C18 high pH chromatography, using homemade EmporeC18 Stage-Tips. Fractions were vacuum-dried and resuspended in 1% (v/v) formic acid, 0.5% DMSO for LC-MS/MS analysis. Peptides were analysed by LC-MS/MS in an Exploris480 mass spectrometer (Thermo) coupled to an UltiMate 3000 RSLCnano LC system (Thermo). Peptides were loaded into a trap column (Acclaim™ PepMap™ 100 C18 LC Columns 5 µm, 20 mm length) for 3 min at a flow rate of 10 µl/min in 0.1% formic acid. Then, peptides were transferred to an EASY-Spray PepMap RSLC C18 column (Thermo) (2 µm, 75 µm x 50 cm) operated at 45 °C and separated using a 60 min effective gradient (Solvent A: 0.1% FA; solvent B: 100% ACN, 0.1% FA) at a flow rate of 250 nL/min. The gradient used was, from 4% to 6% B in 2 min, from 6% to 33% B in 58 min, plus 10 additional min at 98% B. The mass spectrometer was operated in a data-dependent mode, with an automatic switch between MS and MS/MS scans using a top 12 method. (Intensity threshold ≥ 7.5 x 10^4^, dynamic exclusion of 30 sec and excluding charges (+1 and ≥ +8). MS spectra were acquired from 400 to 1500 m/z with a resolution of 60,000 FWHM (200 m/z). Ion peptides were isolated using a 0.7 Th window and fragmented using higher-energy collisional dissociation (HCD) with a stepped normalized collision energy of 29, 31 and 33. MS/MS spectra resolution was set to 60,000 (200 m/z). The ion target values were 3 x 10^6^ for MS (maximum IT of 25 msec) and 1 x 10^5^ for MS/MS (maximum IT set to auto, according to the transient length). For data analysis, Xcalibur raw files were converted into the MGF format through MSConvert (Proteowizard) and used directly as input files for MeroX (Iacobucci et al., 2018). Searches were performed against an ad-hoc protein database containing the sequences of the complexes and a set of randomized decoy sequences generated by the software. The following parameters were set for the searches: maximum number of missed cleavages 3; targeted residues K, S, Y and T; minimum peptide length five amino acids; variable modifications: carbamidomethyl-Cys (mass shift 57.02146 Da), Met-oxidation (mass shift 15.99491 Da); BuUrBu modification fragments: 85.05276 Da and 111.03203 (precision: 10 ppm MS1 and 20 ppm MS2); false discovery rate cut-off: 5%. Finally, each fragmentation spectra were manually inspected and validated.

### Negative staining (NS) electron microscopy (EM)

CCDC6-RET (0.7 μM) was placed onto freshly glow-discharged (GloQube, Quorum) home-made carbon-coated 400 mesh copper electron microscopy (EM) grids. The conditions for glow discharge were: 15 mA, 45“, 0.1 bar. The samples were then incubated for 5 sec at RT. The grids were consecutively placed on top of three distinct 50 μL drops of MilliQ water, striped gently with filter paper for 2 sec and laid on the top of two different 50 μL drops of 1% uranyl acetate and stained for 1 min. Finally, they were stripped gently for 5 sec and air dried. Prepared grids were visualized on a Tecnai 12 transmission EM (Thermo Fisher Scientific) with Lanthanum hexaboride cathode operated at 120 keV. Images were recorded at 61 320 nominal magnifications on a 4kx4k TemCam-F416 CMOS camera (TVIPS), with pixel size of 2.5 Å/pix. For data collection, the open-source software SerialEM (bio3d.colorado.edu/SerialEM) was used under a Massachusetts Institute of Technology (MIT) license. For data processing, open sources software cryoSPARK (cryosparc.com) and Relion were used (Kimanius et al., 2021).

### Small angle X-ray scattering (SAXS)

SAXS experiments were conducted at the beamline B21 of the Diamond Light Source (Didcot, UK)(Cowieson et al., 2020). CCDC6-RETΔCT (40 μL, 5 mg/mL) was injected at room temperature into a Superdex 200 Increase 3.2 column connected to an in-line Agilent 1200 HPLC system. The buffer used for the SEC column was the same protein buffer (20 mM Tris pH 7.5, 500 mM NaCl, 1mM DTT). The eluted samples were exposed for 300 s in 10 s acquisition blocks using an X-ray wavelength of 1 Å, and a sample to detector (Eiger 4M) distance of 3.7 m. The data covered a momentum transfer range of 0.0032 < q < 0.34 Å^-1^. The frames recorded immediately before elution of the sample were subtracted from the protein scattering profiles. The Scåtter software package (www.bioisis.net) was used to analyze data, buffer-subtraction, scaling, merging and checking possible radiation damage of the samples. The R_g_ value was calculated with the Guinier approximation assuming that at very small angles q < 1.3/R_g_. The particle distance distribution, D_max_, was calculated from the scattering pattern with GNOM, and shape estimation was carried out with DAMMIF/DAMMIN, all these programs included in the ATSAS package (Petoukhov et al., 2012). The proteins molecular mass was estimated with GNOM. Interactively generated PDB-based homology models were made using the program COOT by manually adjusting them into the envelope given by SAXS that could match the experimental scattering data. This was computed with the program FoXS (Schneidman-Duhovny et al., 2016)

### Data availability

The mass spectrometry data have been deposited to the ProteomeXchange Consortium via the PRIDE partner repository with the dataset identifier PXD053907 (https://proteomecentral.proteomexchange.org/cgi/GetDataset?ID=PXD053907). The SAXS data was deposited in the small-angle scattering biological data bank (SASBDB) with ID: SASDXXX (https://www.sasbdb.org/data/SASDSXXX). Electron microscopy maps and protein models shown in this study are provided within the submitted manuscript as supporting information.

## Results

### 1. Expression, purification and quality check of a recombinant CCDC6-RET fusion product

We have produced a recombinant CCDC6-RET fusion product (Fig. 1a) composed by fragments of residues 1-101 of CCDC6 and residues 713-1072 of RET using a baculovirus expression system in Sf9 insect cells. Protein purification was achieved by tandem immobilized metal affinity (Ni^2+^) and glutathione-conjugated gravity flow chromatography and in-gel 3C-protease digestion followed by size-exclusion chromatography, if required (Fig. 1b). This is an updated version of a previously published single-step affinity protocol using glutathione-conjugated resin for the purification of recombinant RET kinase and intracellular domains expressed in insect cells using a baculovirus system (Knowles et al., 2006; Plaza-Menacho et al., 2016; Plaza-Menacho et al., 2014a). The tandem double affinity chromatographic step allowed us to obtain highly pure and monodisperse protein with no traces of DNA or nucleotide contamination as indicated by the values of absorbance at 280 and 260 nm and their optimal ratio (Fig. 1c), resulting in a recombinant protein yield of 0.3-0.4 mg per liter of culture. Further quality check over a range of different protein concentrations by mass photometry shows that CCDC6-RET is a 101.2 ± 2.2 kDa dimer in solution (Fig. 1d). These data mirrored the results obtained by SEC-MALS (dimer 97.5 kDa, unpublished data). Binding tightness of RET tyrosine kinase inhibitors BLU-667 (Shehata et al., 2023) and Ponatinib (Mologni et al., 2013) was evaluated by changes in intrinsic fluorescence (IF) upon a temperature gradient (Fig. 1e and f). Binding of both compounds to CCDC6-RET resulted in a shift of the inflection temperature (T_i_) of 10.8 and 20.3 °C, respectively (Fig. 1f). Altogether, these results assured the quality, monodispersion and catalytic integrity of the protein sample prior robust functional and enzymatic characterization. Initial functional characterization showed that CCDC6-RET functions as a highly-active dimer in solution displaying fast autophosphorylation kinetics (Fig. 1g) in agreement with a twenty-fold lower K_M_ value for ATP (22.75 ± 7.5 μM) compared with the RET kinase domain (KD, 429 ± 43 μM) construct (Knowles et al., 2006), see Fig. 1h.

**Figure 1.**
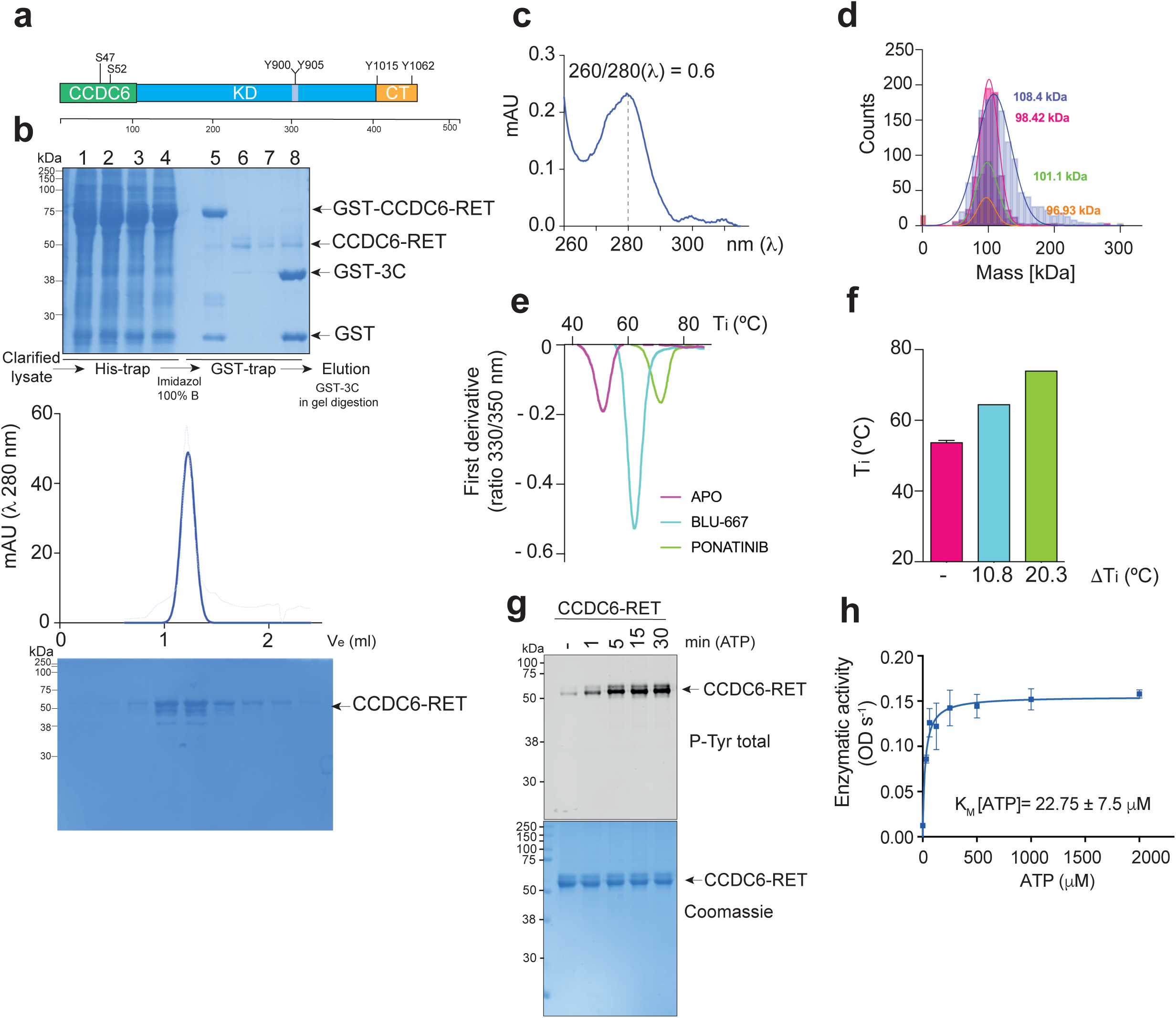
Purification of a recombinant CCDC6-RET fusion product from Sf9 insect cells. **a.** Schematic diagram of the CCDC6-RET fusion product depicting functional domains with the indicated phospho-sites and aa number. **b.** In tandem immobilized metal affinity (Ni^2+^), glutathione-conjugated gravity flow chromatography and in-gel 3C-protease digestion. Indicated fractions were run on an SDS-PAGE and stained with Coomassie: 1-4 elution fractions with 100% buffer B, 5 recombinant protein bound to the glutathione conjugated resin before proteolytic cleavage, 6-7 elution fractions after cleavage with 3C-PreScission protease (GST-tagged, see lane 8), 8 glutathione conjugated resin after proteolytic cleavage. Size exclusion chromatography (sec) using a Superdex 200 3.2/300 column. Indicated fractions were run on an SDS-PAGE and stained with Coomassie (lower panel). **c.** Absorbance spectra (260-310 nm) with max at 280 nm and indicated (260/280 nm) ratio. **d.** Mass photometry profiles (counts vs mass in kDa) of CCDC6-RET at different concentrations in color code: 50 nM (magenta), 25 nM (blue), 10 nM (green), 5 nM (orange). **e.** DSF profile (IF) of CCDC6-RET (2 μM) in apo state and bound to BLU-667 (10 μM) and Ponatinib (10 μM). **f.** T_i_ (°C) and ∇T_i_ (°C) values from f. Data (mean ± SEM) of three experiments, n=3. **g.** WB of samples from a time-course autophosphorylation experiment with CCDC6-RET (1 μM) in the presence of ATP (1 mM) and MgCl_2_ (2 mM) for 0–30 min using a total anti-phospho-tyrosine antibody. The total amount of protein was visualized by Coomassie staining. **h.** Enzymatic assay performed with CCDC6-RET (1 μM) incubated with increasing concentrations of ATP at a fixed concentration (1.35 mM) of a c-Abl derived peptide (EAIYAAPFAKKK). Data represent the mean ± SEM of two experiments in duplicate (n = 4). Enzymatic activity (ODs^−1^) versus ATP concentration and K_M_ values are depicted.

### 2. Mass spectrometric characterization of CCDC6-RET autophosphorylation reveals dual-specificity kinase activity

Next, we applied high-resolution time-resolved liquid chromatography tandem mass spectrometry (LC-MS/MS) to monitor the dynamic changes of the autophosphorylation reaction on CCDC6-RET *in vitro*. Fast autophosphorylation kinetics were observed in most of the identified phospho-sites, which recapitulated the results from the WB data using phospho-specific antibodies (Fig. 2a). A total of twenty phospho-sites were mapped, of which seventeen were *the facto* autophosphorylation sites as indicated by their time-dependent saturation kinetics during the reaction (Fig. 2b and c), see also supplemental Table 1 and supplemental data file 1. CCDC6-RET display dual-specificity kinase activity as it is able to autophosphorylate on both serine and tyrosine residues. CCDC6-RET autophosphorylates not only on sites located on the catalytic core of the fusion but also on discrete sites of the N-terminal and coiled-coil region of CCDC6, some of which were not previously described. We identified among other several phospho-sites on the amino terminal and coiled-coil region of the gene fusion product: Ser 46, Ser 52, Tyr 80 and Ser 98 (Fig. 2c, d). Among these, Ser 6 and Tyr 80 are novel phosphosites on CCDC6 (Fig. 2c, d). Mutating Ser 46, Ser 52 and both residues simultaneously to alanine does not cause any defects on the phospho-tyrosine activity as indicated by the autophosphorylation reaction (Supplementary Fig. 1a-c). In the same line, a CCDC6-RET fusion with a Y80F or a S98A mutation appeared to be catalytic competent (Supplementary Fig. 1d and e). Autophosphorylation sites on the catalytic core of the RET fusion were also identified (Supplementary Table 1 and supplemental data file 1). These phospho-sites are located at different structural and functional motifs of the RET catalytic domain e.g. Tyr 806 and Tyr 809 (hinge), Tyr 826 (kinase insert), Ser 896, Tyr 900, Ser 904, Ser 905, Tyr 905, Ser 909 (activation loop) and Tyr 1015, Tyr 1029, Ser 1021, Ser 1034 and Tyr 1062 (c-terminal segment), see also Fig. 2c. Some of these phospho-sites showed high levels of phosphorylation in the basal state which decreases once the autophosphorylation reaction evolved during time. This is the case for activation loop Tyr 900 and c-terminal Ser 1034, both phospho-sites are located within peptides containing in addition another bona fide autophosphorylated residues whose levels increase one order of magnitude during the reaction, see on Fig. 2f the example of activation loop tyrosine 900 and 905. In other cases, phospho-sites appeared only in doubly or triply phosphorylated peptides (e.g. Ser 896 and Ser 1021). A complete list of the CCDC6-RET phospho sites identified in at least two out of three independent experiment is shown on Supplementary table 1. Next, we targeted c-terminal phospho-sites to test whether they are required for catalytic activity. The mutation of residue Tyr 1015 to phenylalanine (Y1015F) did not cause any detrimental effect on the phospho-tyrosine activity as seen in the time course autophosphorylation assays (Supplementary Fig. 1g). The same applied to a Y1062F and S1034A mutants (Supplementary Fig. 1g and h). As negative controls we used adenosine mono-phosphate (AMP), and bi-phosphate (ADP) nucleotides (Supplementary Fig. 2a and b), and we found a significant increase on CCDC6-RET phospho-tyrosine activity in the presence of ADP and not by AMP. These unexpected results were a clear sign of ADP-dependency by the CCDC6-RET fusion product, which we explored further (see next result section).

**Figure 2.**
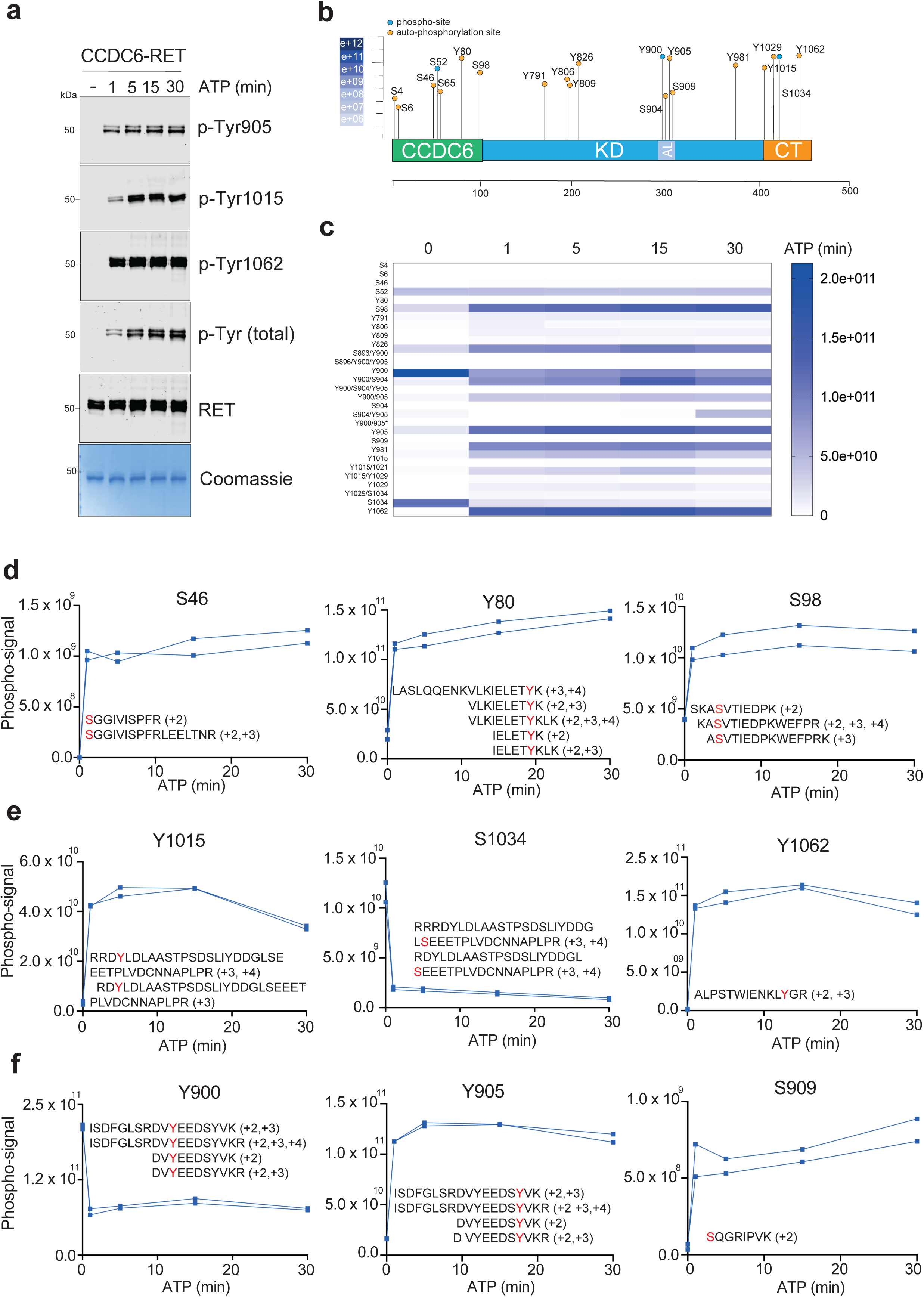
LC-MS/MS characterization of CCDC6-RET autophosphorylation reveals dual-specificity kinase activity. **a.** WB of samples from a time-course autophosphorylation experiment with CCDC6-RET (1 μM) in the presence of ATP (1 mM) and MgCl_2_ (2 mM) for 0–30 min using the indicated total and phospho-specific antibodies. The total amount of protein was visualized by Coomassie staining. **b.** Schematic diagram of the CCDC6-RET fusion product with the identified phospho-sites and aa number (note hereafter that c-terminal sites of the fusion follow the RET aa numbering) that resulted from the LC/MS-MS analyses. **c.** Heat map (intensity, white-blue scale arbitrary units) representation of the phospho-proteomic quantification from the CCDC6-RET autophosphorylation reaction in vitro (0-30 min, ATP) identified by LC/MS-MS. **d-f**. Phosphorylation kinetics (relative abundance from time zero, unstimulated) of the indicated CCDC6-RET phospho-sites (highlighted in red, with the different tryptic peptides and charges) measured by LC-MS/MS from c. Data shown are two technical replicates of a single experiment. Data is representative of two independent experiments.

### 3. CCDC6-RET is a dual ATP- and ADP-dependent kinase

Next, we performed a direct functional comparison of CCDC6-RET with a RET 713-1072 aa fragment, the catalytic core of the fusion product, as a control (Fig. 3a). First, we performed enzymatic assays to calculate the catalytic rates and enzymatic constants for ATP using an excess of peptide as an exogenous substrate (Fig. 3b). Under these experimental conditions, CCDC6-RET was a more active enzyme than the catalytic core (aa 713-1072) of the RET fusion as indicated by a six-fold increase in the catalytic efficiency constant (k_cat_/K_M_,) by the oncogenic chimera (Fig. 3c). We next performed *in vitro* time course autophosphorylation assays (0–30 min) using ATP (1 mM), MgCl_2_ (2 mM) with 1 μM final protein concentration. Autophosphorylation was monitored by Western blotting (WB) using RET total and phospho-specific against Tyr 905, Tyr 1015, Tyr 1062 and also total phospho-tyrosine antibodies (Fig. 3d). Fast CCDC6-RET autophosphorylation kinetics reaching saturating levels at early time points (1 min) contrasted with slower rates of autophosphorylation by the 713-1072 construct, see e.g. Tyr 905 phosphorylation (Fig. 3d). We also evaluated the binding of ATP and ADP to both constructs respectively by monitoring changes on IF (Fig. 3 h-j). Titration experiments using a range of increasing concentrations of both nucleotides allowed us to calculate an apparent K_D_ for ATP (186.8 ± 38.3 μM) and ADP (1057.0 ± 157.3 μM) respectively (Fig. 3j). These results together with the MS data (Fig. 3g and Supplemental Fig. 2a and b) prompted us to explore further whether CCDC6-RET is a dual ATP- and ADP-dependent kinase that can use in addition to ATP, ADP as a phosphoryl donor. We found that in contrast to a 713-1072 construct, CCDC6-RET efficiently auto-phosphorylates in the presence of ADP (Fig. 3e-g), although with slower rates than with ATP, in accordance with the affinity values obtained (Fig. 3j). Together these data demonstrate that CCDC6-RET is a highly active dimer in solution with fast kinetics of autophosphorylation that functions as a dual ATP- and ADP-dependent kinase able to bind to and use both ATP and ADP as phosphoryl donors.

**Figure 3.**
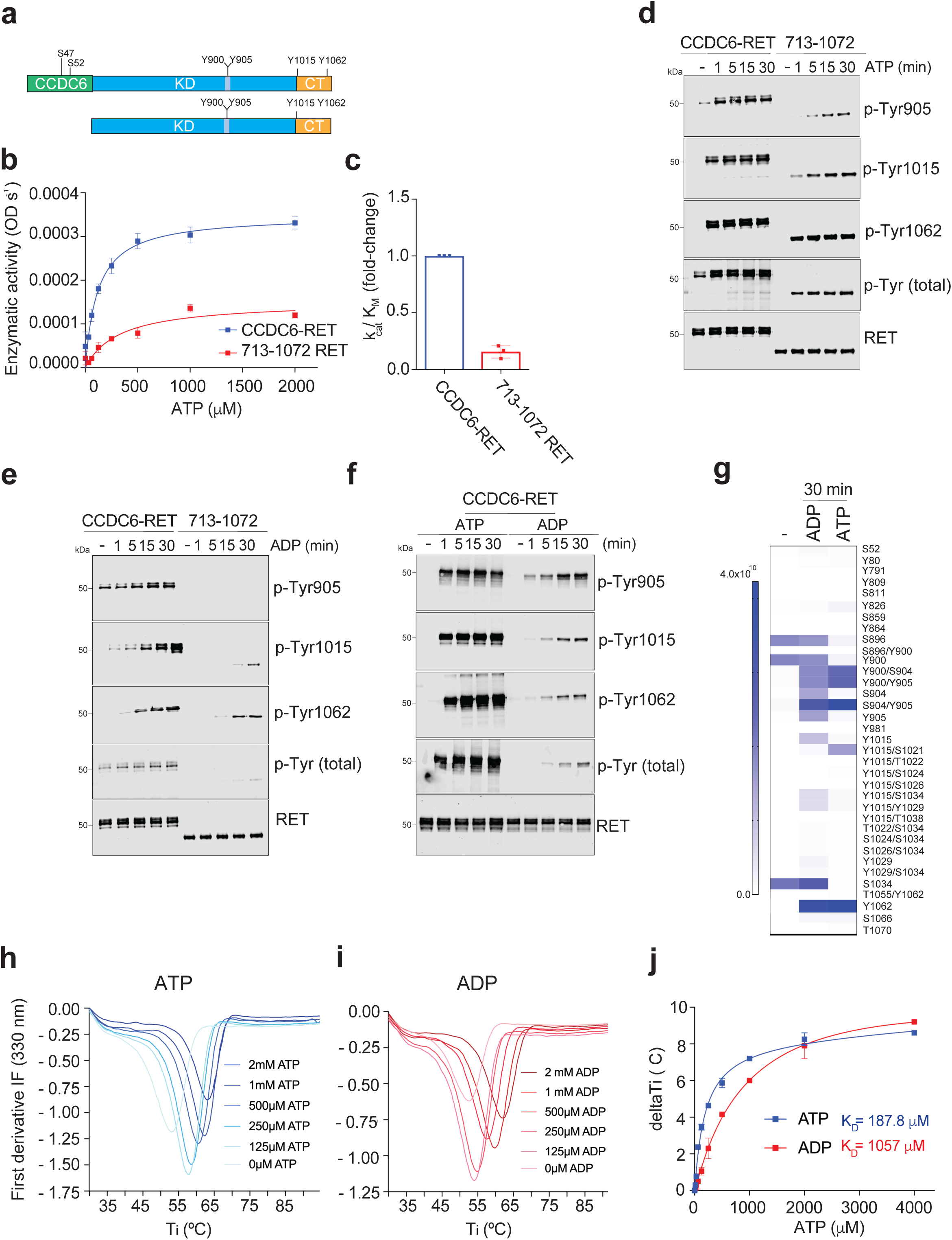
CCDC6-RET is an ADP-dependent kinase. **a.** Schematic diagram of the CCDC6-RET fusion product and RET catalytic core depicting functional domains with the indicated phospho-sites and aa number. **b.** Enzymatic assay performed with CCDC6-RET and RET aa 713-1072 (0.3 μM) incubated with increasing concentrations of ATP at a fixed concentration (1.35 mM) of a c-Abl derived peptide (EAIYAAPFAKKK). Data represent the mean ± SEM of two experiments in duplicate (n = 4). **c.** Catalytic efficiency constant (K_M_/k_cat_) fold-change versus control (i.e. RET aa 713-1072) from b. **d**-**f**. WB of samples from an in vitro time-course autophosphorylation experiment with CCDC6-RET and RET aa 713-1072 (1 μM) in the presence of ATP or ADP (1 mM) and MgCl_2_ (2 mM) for 0–30 min using the indicated antibodies. **g**. Heat map (white-blue scale) representation of the phospho-proteomic quantification captured by LC/MS-MS from the CCDC6-RET autophosphorylation reaction in vitro 0 and 30 min after ATP (1 mM) and ADP (1 mM) treatment in presence of 2 mM MgCl_2_. **h, i**. Representatives DSF profiles (IF, emission 330 nm, first derivative) of CCDC6-RET upon titration with increasing concentrations of nucleotides (ATP and ADP) showing the transitions of the inflexion temperature (T_i_) **j**. Apparent K_D_ values for ATP and ADP (μM). Increment of the T_i_ (∇ T_i_, °C) versus the concentration of nucleotide (μM). Data shown is the mean value ± SEM of three experiments (n = 3).

### 4. CCDC6-RET lacking the c-terminal segment is more catalytic efficient

RET has a highly flexible c-terminal segment that lacks any secondary structural elements but contains critical tyrosine residues required for downstream signalling. To assess if this segment, which is *per se* an intrinsically disordered region (IDR) (van der Lee et al., 2014), plays a role in the regulation of the catalytic activity of the fusion product, we generated a construct lacking the c-terminal segment (missing aa 1014-1072) called ΔCT, see Fig. 4a. Next, we evaluated the catalytic activity of this construct versus the wild-type fusion in enzyme kinetics experiments. First, we used a peptide substrate derived from the RET activation loop (DVYEEDSYVK) and found that the ΔCT construct was more efficient in phosphorylating the activation loop derived peptide (Fig. 4b and c) as shown by the higher catalytic efficiency constant (k_cat_/K_M_) for ATP. The same results were obtained when using an c-Abl-derived peptide (Fig. 4d and e). From these experiments we conclude that: i) the c-terminal segment of the RET fusion product is not required for the catalytic activity, and ii) the c-terminal segment must somehow restrict sterically the access of the substrate into the active site, by contacting key secondary structural elements and functional motifs that define or are within the vicinity of the active cleft. Therefore, a ΔCT construct appears to be a more catalytic efficient enzyme. In fact, crosslinking MS data on CCDC6-RET demonstrate that the c-terminal segment appears to be a highly dynamic node that interacts with distinct secondary structural elements and functional motifs of the kinase domain (i.e. GRL, αC helix and hinge), as indicated by the high number of crosslinked peptides converging on residues surrounding Tyr 1062, mainly on K1060, see figure 4f. Altogether these results suggest that the c-terminal segment of CCDC6-RET can sterically restrict the access into the active site, hence acting as a repressive element for the phosphorylation of exogenous susbtrates.

**Figure 4.**
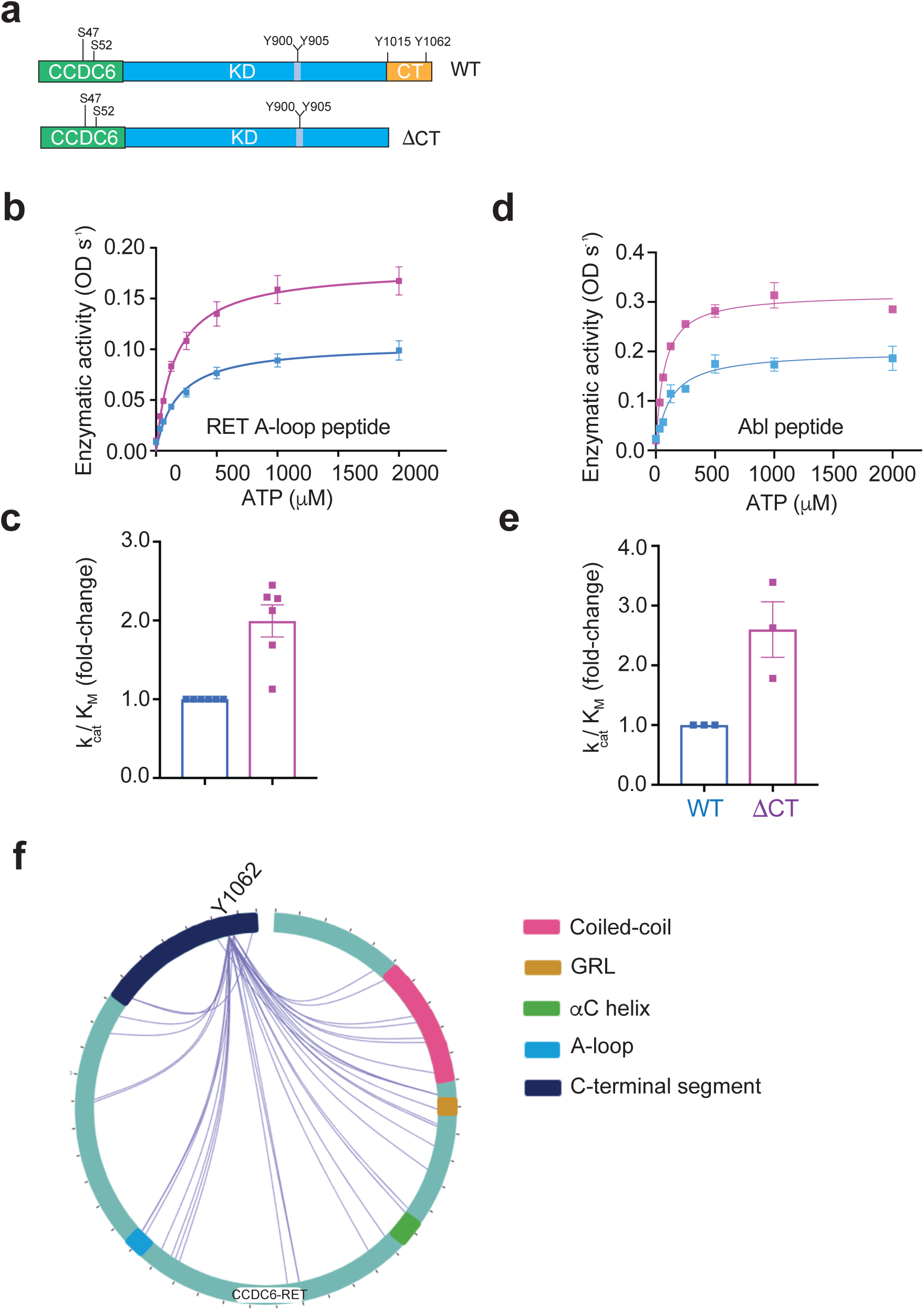
CCDC6-RET lacking the c-terminal segment is more catalytic efficient. **a.** Schematic diagram of the CCDC6-RET fusion products, a wild-type and a c-terminal deleted variant (∇CT, lacking aa 1013-1072) depicting functional domains with the indicated phospho-sites. **b.** Enzymatic assay performed with CCDC6-RET and CCDC6-RET ∇CT (0.2-0.3 μM) incubated with increasing concentrations of ATP at a fixed concentration (2.9 mM) of a RET activation-loop derived peptide (DVYEEDSYVK). Data enzymatic activity (ODs^-1^ x10^-3^) represent the mean ± SEM of three experiments in duplicate (n = 6). **c.** Catalytic efficiency constant (K_M_/k_cat_), fold-change versus control (CCDC6-RET) from b. **d.** Enzymatic assay performed with CCDC6-RET and CCDC6-RET ∇CT (0.2-0.3 μM) incubated with increasing concentrations of ATP at a fixed concentration (1.35 mM) of the c-Abl derived peptide. Data enzymatic activity (ODs^-1^ x10^-3^) represent the mean ± SEM of three experiments in duplicate (n = 6). **e.** Catalytic efficiency constant (K_M_/k_cat_), fold-change versus control (CCDC6-RET) from d. **f.** Circular diagram representation of de cross-linking MS data showing intramolecular crosslinked peptides converging on a c-terminal node around Tyr 1062.

### 5. Catalytic dependency on activation loop Tyr 905 by a CCDC6-RET fusion lacking the c-terminal segment

It is known that the activation loop of protein kinases in the unphosphorylated state can function as an autoinhibitory module by sterically blocking substrate access into the active site (Hubbard et al., 1994). Based on our previous results indicating that the c-terminal segment can also restrict the access of the susbtrate into the catalytic cleft (see Fig. 4b-f), we aimed to explore further a potential crosstalk between the activation loop and the c-terminal segments in the mechanism of CCDC6-RET autoactivation and signalling. For this purpose, we generated a series of activation loop mutants in full-length and c-terminal deleted constructs (ΔCT). First, we produced single and double activation loop tyrosine mutants in a full-length construct (Y900F, Y905F and Y900/905F) and evaluated their catalytic activity in time course autophosphorylation experiments (Fig. 5a). None of the evaluated mutants had a detrimental impact on the phospho-tyrosine activity compared with the WT control (Fig. 5a). Of note, a full-length CCDC6-RET Y900F mutant construct resulted consistently in very low yield of soluble protein after purification. Although it was catalytically competent (Supplementary Fig. 3), the very low amount of recovered material made it difficult to work at comparable levels due to its short life and that it was not possible to concentrate it to reach workable concentrations. Intriguingly, such detrimental effect on protein yield was rescued by the concomitant mutation on Tyr 905 in the case of a double activation loop mutant (Y900/905F), see Fig. 5a. On the contrary, when we evaluated the function of the complete series of activation loop mutants in a c-terminal deleted context (ΔCT), we observed that the Y900F mutant was efficiently expressed to the same levels as the WT protein. Interestingly, we also observed a catalytic dependency on activation loop Tyr 905 in the ΔCT constructs (Fig. 5b). In particular, we found that in autophosphorylation and enzyme kinetics experiments, where we evaluated the K_M_ for ATP in the presence of an excess of peptide susbtrate, a Y905F and double Y900/905F mutants have a loss of function effect, contrary to the Y900F, that was catalytic competent. Hence, these data uncovered a catalytic dependency on Tyr 905 by the CCDC6-RET fusion product when the c-terminal in not available. Furthermore, these data also indicate that Tyr 900 appears to be critically involved in the unfolding and processing of the intact protein (i.e. in the presence of the c-terminal segment) being both discrete and mutually exclusive events. It is evident from these data that a key steric interaction between (phosphorylated) activation loop Tyr 900 and c-terminal elements are required for the proper folding of the oncogenic chimera. This interaction appears to be complemented by a concomitant mechanism, where Tyr 905 appears as a key molecular determinant implicated in the transduction of the oncogenic signal once the c-terminal segment is not available (e.g. once it is engaged in the assembly of a signalling complex after autophosphorylation). From these data, we conclude that the CCDC6-RET activation loop phospho-sites control both the catalytic activity and the unfolding of the fusion protein by a mutually exclusive c-terminal segment dependent mechanism.

**Figure 5.**
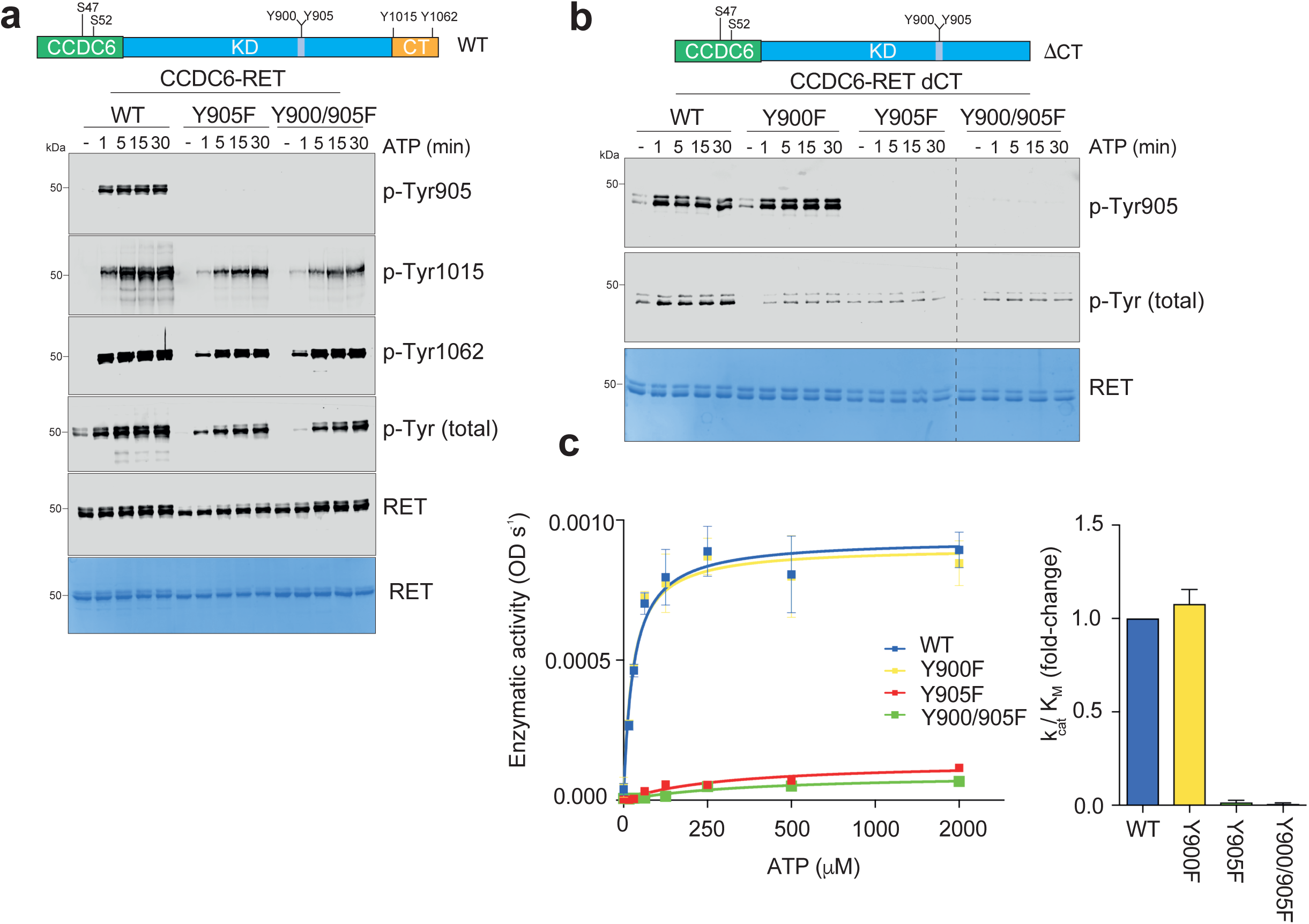
Activation loop phospho-sites controls both the catalytic activity. **a.** Schematic diagram of the CCDC6-RET fusion product depicting functional domains with the indicated phospho-sites and aa number. WB of samples from a time-course autophosphorylation experiment with CCDC6-RET WT and activation loop mutants Y905 and Y900/905F (1μM) in the presence of ATP or ADP (1 mM) and MgCl_2_ (2 mM) for 0–30 min using the indicated antibodies. **b.** Schematic diagram of the CCDC6-RET ∇CT fusion product depicting functional domains with the indicated phospho-sites and aa number. WB of samples from a time-course autophosphorylation experiment with CCDC6-RET ∇CT WT and activation loop mutants Y900F, Y905 and Y900/905F (1μM) in the presence of ATP or ADP (1 mM) and MgCl_2_ (2 mM) for 0–30 min using the indicated antibodies. **c.** Enzymatic assay performed with CCDC6-RET ∇CT, WT and activation loop mutants Y900F, Y905 and Y900/905F (1μM) incubated with increasing concentrations of ATP at a fixed concentration (1.35 mM) of a c-Abl derived peptide (EAIYAAPFAKKK). Data represent the mean ± SEM of two experiments in duplicate (n = 4). Catalytic efficiency constant (K_M_/k_cat_) fold-change versus control (WT).

### 6. *In silico* reconstruction of a CCDC6-RET homodimer for single particle analyses

We generated a high-confidence structural reconstruction of a CCDC6-RET homodimer *in silico* for electron microscopy (EM) single particle analyses. First, we used the Waggawagga coiled-coil prediction server (Simm et al., 2015). Coiled-coil predictions are based on the identification of periodic heptad repeats, which can be represented by a net- or helical wheel-diagrams. This bioinformatic tool enables the discrimination between coiled-coil-domains and single α-helices (SAH). In the case of CCDC6-RET there is a high confidence parallel homodimer coiled-coil interface from residues 56-101 of each protomer (Fig. 6a and d), as indicated by both the coiled-coil probability marcoil (p-value 100) and very low SAH-score (0.0333). The coiled-coil inner hydrophobic core was clearly defined by the heptad repeats (Fig. 6b and c), and several salt-bridge interactions (strong) involving Glu 58 and Arg 62, Lys 74 and Glu 78, and Lys 85 and Glu 89. These contacts were complemented by middle strength interactions involving Glu 78 and Lys 81 and Glu 89 and Arg 92, and weak coordination between Asp 93 and Lyś96, as depicted in the helical wheel- and heptad-net-diagrams (Fig. 6b and c). We visualized in 3D a parallel homodimeric coiled-coil interface (Fig. 6d) using CCbuilder 2.0 (Wood and Woolfson, 2018). In an attempt to disrupt the dimerization interface, we generated a series of mutants at key positions within the coiled-coil, for example Cys 84 (C84A) that can form a disulfide bridge with its counterpart (Fig. 6d), and Ser 98 (S98A) which we found was an unexpected autophosphorylation site, see supplementary table 1. None of these mutants were able to dissociate the dimer as indicated by the elution volumes observed in SEC analyses compared with the WT (data not shown). These coiled-coil mutants did not affect the activity of the fusion products as detected in autophosphorylation experiments (Supplementary Fig. 1e and f), suggesting a tight dimerization constant by the interface. Next, we applied artificial intelligence to obtain a high-confidence 3D-predictive model of a monomeric CCDC6-RET fusion protomer using AlphaFold (AF id: J7M8C2). By the superimposition of two monomers of the predicted fusion product onto the dimeric coiled-coil interface (Fig. 6d) we generated a two-fold rotational symmetry dimer (Fig. 6e) in which the catalytic domains are facing away symmetrically from the coiled-coil stem. The high confidence and feasibility of the predictive 3D-model was supported by careful inspection of key structural features, see Supplemental Fig. 4 a-c: i) the correct position and configuration of secondary structural elements and functional motifs of an active kinase defined overall by the correct alignment of the regulatory (R-) and catalytic (C-) spines and DFG-in configuration (Plaza-Menacho et al., 2016); ii) the hydrophobic patch formed by contacts between the αC helix (i.e. PIF-like pocket) and the coiled-coil stem; iii) analogy to the domain arrangement found in bacterial histidine kinases where the catalytic domain is place to the left of the symmetry plane of the coiled-coil interface defined in those cases by a four helix bundle instead (Bhate et al., 2015), and iv) in good agreement with two different solutions provided by AlphaFold multimer that could represent different configuration states in light of the specific arrangements of the catalytic domains with respect to the coiled-coil interface (Supplemental Fig. 4 e and f).

**Figure 6.**
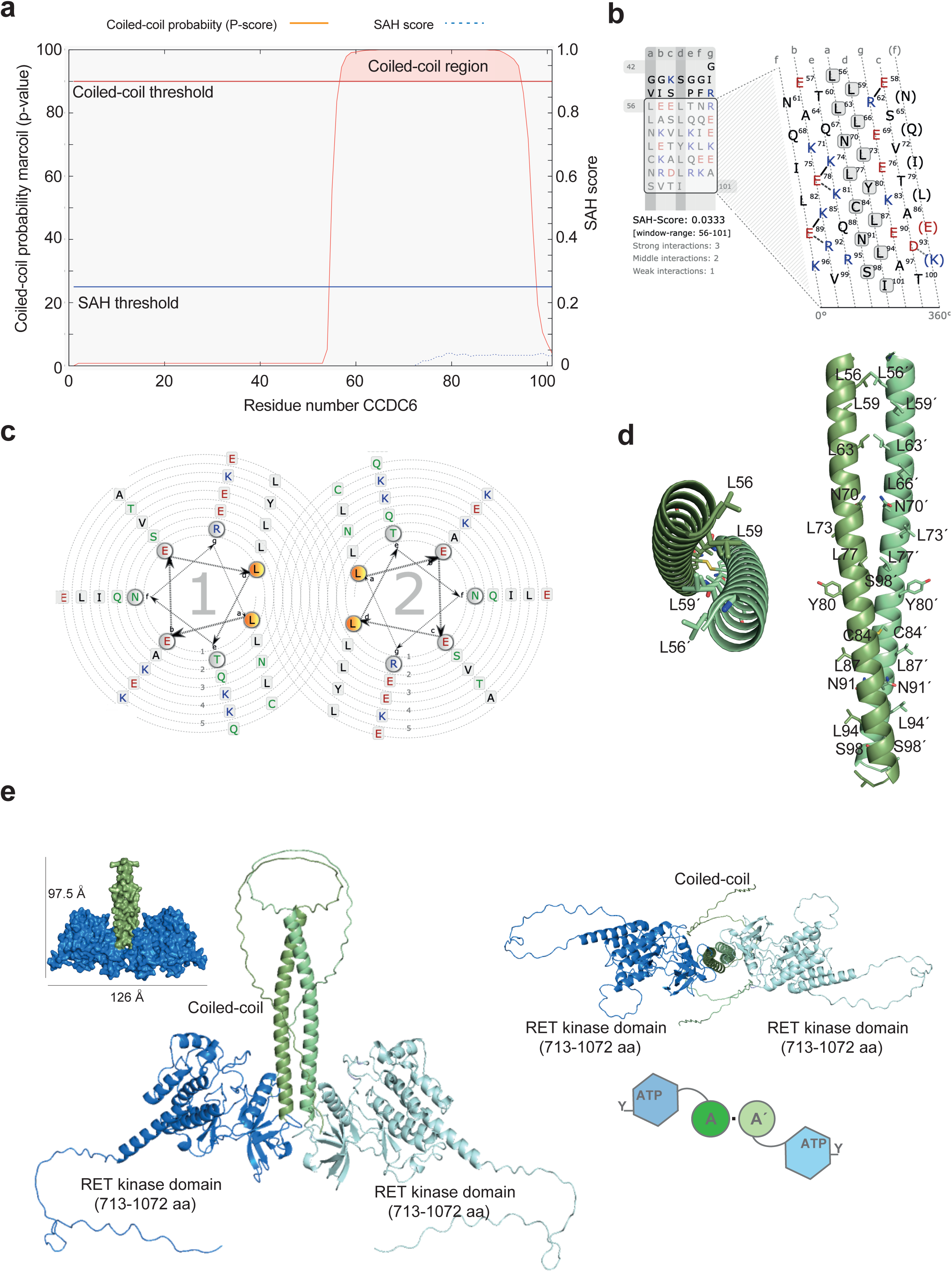
In silico reconstruction of a 3D-model of a CCDC6-RET homodimer. **a.** Coiled-coil probability (p-score) and single alpha helix (SAM) score for marcoil prediction of the n-terminal sequence of CCDC6-RET. **b.** Identification of the heptad repeats depicted in the form of a net-diagram for the coiled-coil region (aa 54-101) of CCDC6-RET. **c.** Heptad repeats represented in a helical wheel view, from a and b. **d.** 3D-visualization (cartoon representation) of the coiled-coil region (54-101) of CCDC6-RET with the side chains of the represented residues in sticks. Frontal and lateral views are depicted with indicated side chain residues. **e.** A high-confidence 3D-predictive model of a monomeric CCDC6-RET fusion protomer comprising the short c-terminal isoform of RET using Alpha-Fold (AF id: J7M8C2). Superimposition of two monomers of the predicted fusion product onto the parallel homodimeric coiled-coil interface generates a two-fold rotational symmetry dimer in which the catalytic domains are facing away symmetrically from the coiled-coil stem. 3D-visualization of the coiled-coil region of the CCDC6-RET dimer. Frontal and upper views are depicted and schematic diagram of the domain organization showing the 2-fold rotational symmetry arrangements domains (top view).

### 7. Architecture of a CCDC6-RET dimer by EM single particle analyses

CCDC6-RET (0.7-1 μM) was placed on home-made carbon-coated 400 mesh copper grids, negatively stained, and analyzed by single-particle methods to obtain an empiric 3D reconstruction (Fig. 7a). Reference-free classification performed with Relion (Kimanius et al., 2021) gave a range of 2D-class averages from which a parallel homodimeric coiled-coil interface linked to two globular domains in different orientations with apparent 2-fold symmetry were extracted (Fig. 7b). Single particle images of a CCDC6-RET dimer revealed a range of different orientations and a good level of detail in the overall architecture that was in good agreement with a dimeric arrangement of the functional domains; and the size and shape of the in silico reconstructed model, where IDRs were not included (126 Å in its longer dimension and 97.5 Å in its shorter dimension), see Fig. 7 b and c. The reference free 2D-classes concur well with the retro-projections of the reconstituted 3D-model adopting an inverted Y shape, with a thinner stem (coiled-coil interface) in the middle and two globular arms segregated from the stem facing at opposite almost perpendicular directions (Supplemental Fig. 5a); similar 2D-classes were obtained with cryoSPARK (Punjani et al., 2017) see Supplemental Fig. 5b. A starting 3D model was built from such 2D-classes assumed to correspond to almost orthogonal orientations of the particle on the grid. This initial model was further refined to produce the final EM map (Fig. 7d and e, and supplementary Fig. 5c). The reliability of the map is attested by the good agreement between classes and reprojections, the even distribution of Euler angles (supplementary Fig. 5d), our 3D-reconstructed and dimeric models from the AlphaFold multimer (see Supplemental Fig. 4). The 3D-map confirms the dimeric and twisted arrangement of the globular catalytic domains towards the left of the 2-fold rotational axis from the coiled-coil stem. This comes in line with the catalytic domain configuration seen in bacterial histidine kinases (Bhate et al., 2015), where the catalytic core of the DHp-CA module is placed to the left of helix b. When the density map is contoured at a threshold chosen to encompass a molecular weight of 110 kDa (calculated molecular weight of CCDC6-RET), the map generally has good connections between adjoining domains but lacks density for the whole coiled-coil interface and N-terminal part of the fusion.

**Figure 7.**
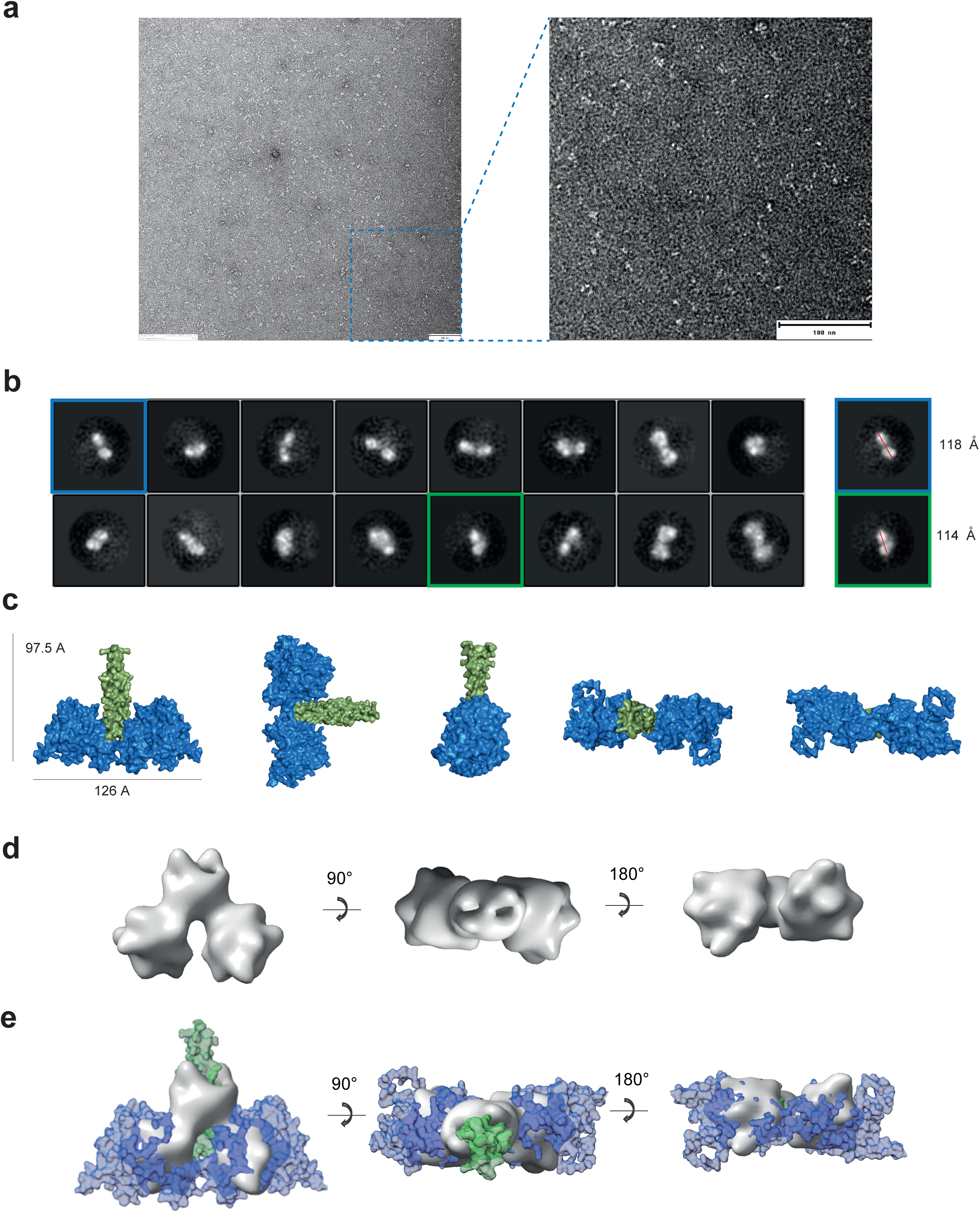
Architecture of a CCDC6-RET dimer by EM single particle analyses. **a.** Micrograph of a representative NS grid view, scale bar 100 nm. Magnified view on the right panel. **b.** Single particle analyses reference free 2D-classification of CCDC6-RET, showing representative single particle views and size. **c.** Orthogonal views of an in-silico 3D-reconstructed CCDC6-RET dimer showing particle size. Note, amino- and carboxy-terminal CCDC6-RET IDRs regions are not included. Coiled-coil region represented in green and catalytic domains in blue color code. **d.** 3D-maps (surface representation) of CCDC6-RET, in different orthogonal orientations (frontal, upper and bottom views). **e.** Superimpositions of 3D-maps (surface representation) from d, onto the in-silico 3D-reconstructed CCDC6-RET dimer from c, in different orthogonal orientations (frontal, upper and bottom views).

To obtain further information on the molecular shape of a CCDC6-RET homodimer in solution, low-angle X-ray scattering (SAXS) data of a CCDC6-RET-ΔCT construct were collected (Supplementary Fig. S6a). Pair distance distributions exhibited fine features consistent with a multidomain protein sample. Ab initio envelopes were generated for CCDC6-RET-ΔCT using DAMAVER consistent with the pair distance distributions showing an elongated shape at the root of the coiled-coil and a more globular architecture at the catalytic portion of the fusion resulting in an inverted T-shape object (Supplementary Fig. S6b and c) with a radius of gyration (Rg) of 55.07 ± 0.18 Å and D_max_ of 195 Å. These measurements are in accordance with the reference-free 2D-particles and 3D-envelop obtained by electron microscopy and also the 3D-in silico reconstructed particles (see Fig. 6 and 7). Of note, the CCDC6-RET-ΔCT construct generated an extra volume or “bump” adjacent to one of the two catalytic domains of the fusion, which was tentatively attributed to the dynamic motion of both catalytic domains, as we can fit in another catalytic domain molecule within the extra volume. The CCDC6-RET-ΔCT structure was modeled using an Alpha-Fold (AF id: J7M8C2) of a monomeric chimera without the amino- and carboxy-terminal IDRs. The initial 54 amino acids were then then manually built into the SAXS envelope (Supplemental Fig. 6 d). The fitting of the theoretical curves derived from this CCDC6-RET-ΔCT model against the experimental curves gave a χ^2^ value of 2.2, indicating a good quality of the data, see supplemental table 3. The high-quality SAXS data were therefore sufficient to generate a preliminary model for a CCDC6-RET dimer assembly, defining clear pairwise interdomain angles of approximate 90° between the coiled-coil interface and the catalytic domains and a potential solution for the extra volume observed in the SAXS envelop (Supplemental Fig. 6d). The coiled-coil and catalytic domains angle appears to be in the range observed for the DHp-and CA domains in bacterial histidine kinases (Bhate et al., 2015; Dubey et al., 2020; Dubey et al., 2016). The SAXS envelops fits well all the in silico generated CCDC6-RET assemblies and the 3D-map from the EM single particle analyses, capturing the dynamic transitions between the different assemblies’ states and the dynamic motion of the catalytic domains from the coiled-coil stem. These data also suggest some degree of asymmetry between catalytic domains and the dimer interface (Supplemental Fig. 6d).

### 8. A mechanism *in cis* drives CCDC6-RET autophosphorylation

Modes of kinase autophosphorylation are based on the dimerization dependence of the reaction. The observation of a concentration-dependent increase in kinase activity is interpreted as a bi-molecular reaction mechanism in trans (Reinhardt and Leonard, 2023). Otherwise, autophosphorylation reactions that are not influenced by dimerization are not subject to any intermolecular regulation, hence they proceed *in cis* (Reinhardt and Leonard, 2023). In order to investigate these two components in the mechanisms of CCDC6-RET autoactivation, we performed a series of time-course autophosphorylation reactions *in vitro* at increasing enzyme concentrations (2.5, 1 and 0.25 μM). WBs analyses using total phospho-tyrosine and RET phospho-specific antibodies showed a clear concentration independent effect on the speed of the reaction, indicative of a reaction *in cis* (Fig. 8a). As a control, we performed the same experiment using the catalytic core (RET aa 713-1072) of the fusion protein instead (Fig. 8a). In this case, as previously seen for the complete intracellular domain (ICD) (Plaza-Menacho et al., 2014a), RET autophosphorylation kinetics showed a marked enzyme concentration dependency, which was indicative of an intermolecular process (i.e. autophosphorylation *in trans*). Next, we evaluated the enzymatic activity of CCDC6-RET at incremental enzyme concentrations (6, 3 and 1 μM) in an in vitro kinase assay using increasing ATP concentrations (Fig. 8b). Again, a lack of enzyme concentration effect on the catalytic rates of CCDC6-RET was observed, as indicated by the unchanged catalytic constant (k_cat_) see Fig. 8b. This was in line with the previous set of results (Fig. 8a) and supports the notion of a unimolecular reaction, in which CCDC6-RET fusion autophosphorylation is driven by a mechanism *in cis*. A gold standard method to discriminate between *cis*- and *trans*-autophosphorylation is to evaluate if an active kinase can phosphorylate intermolecularly an inactive susbtrate surrogate that mimics its own kinase domain (Cuesta-Hernandez et al., 2023; Reinhardt and Leonard, 2023). This approach can also be performed in the context of both, a native or an obligate, artificial heterodimer. Time course experiments of CCDC6-RET autophosphorylation in the presence of an intact susbtrate surrogate were performed. In this case an amino- and carboxy-terminal deleted and HRD catalytic inactive mutant (∇CCDC6-RET-HRA) that is catalytic inactive was used as a susbtrate surrogate (Fig. 8c). WBs data using a total phospho-tyrosine and RET activation loop phospho-specific antibodies revealed that CCDC6-RET display a very strong tendency to autophosphorylate to itself rather than to the inactive kinase susbtrate (Fig. 8c), which is indicative of *cis*-autophosphorylation. Note that the levels and kinetics of CCDC6-RET autophosphorylation are unchanged in the presence of the susbtrate molecule, supporting our interpretation. In fact, when evaluating in the same experimental setting another protein kinase that it able to autophosphorylate on the activation loop *in trans* (e.g. c-Src) we observed a totally opposite output (Fig. 8c). In this case, we observed a very rapid and significant susbtrate surrogate phosphorylation, which is accompanied by a slow-down in the kinetics of autophosphorylation of the active kinase in the presence of the substrate molecule (Fig. 8c).

**Figure 8.**
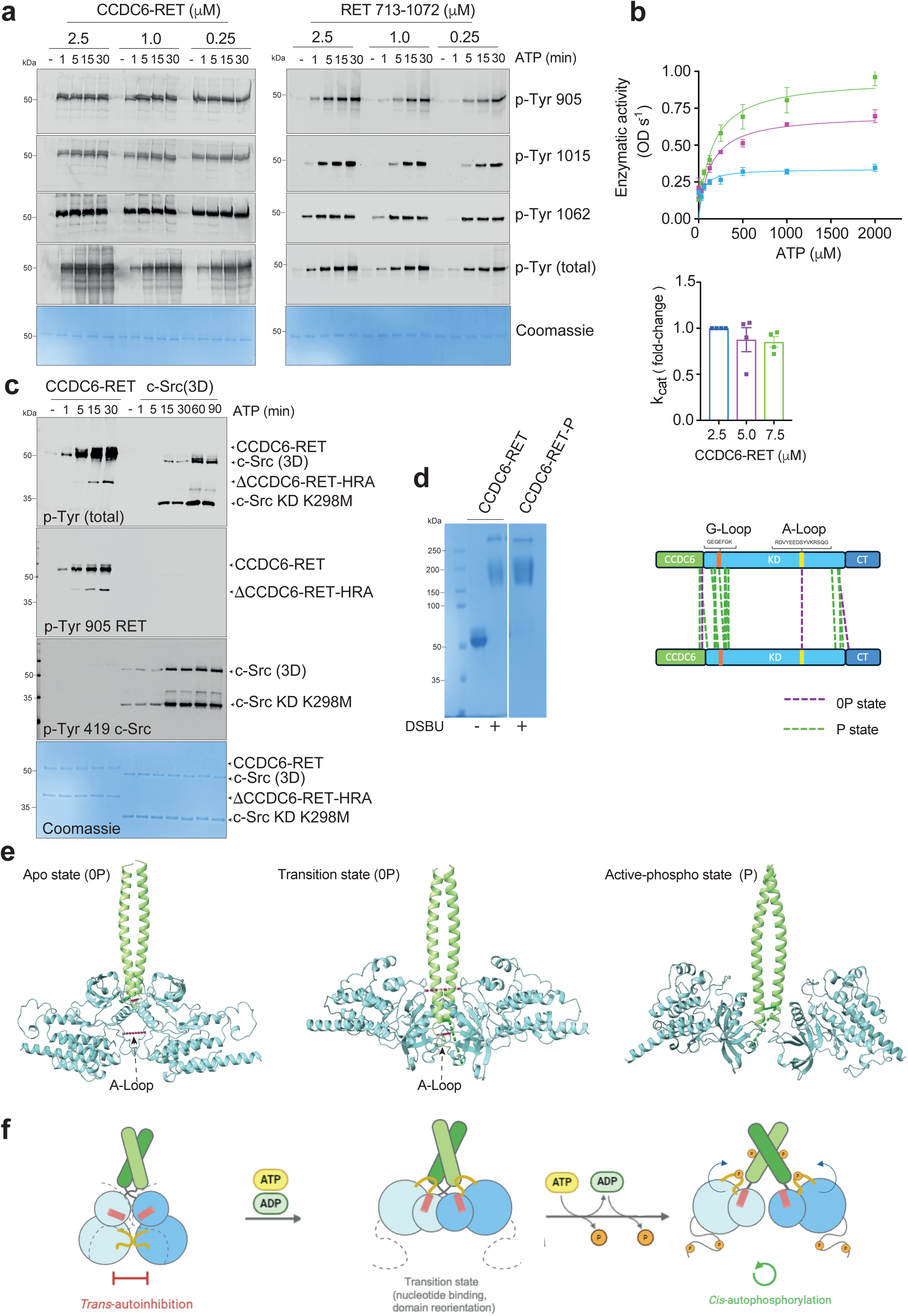
A *cis*-mechanism for CCDC6-RET autophosphorylation. **a.** WB of samples from a time-course autophosphorylation experiment with CCDC6-RET and a RET catalytic core fragment (aa 713-1072 fragment) at increasing (0.25–2.5 μM) enzyme concentrations in the presence of ATP (1 mM) and MgCl_2_ (2 mM) for 0–30 min using total phospho-tyrosine and the indicated RET phospho-specific antibodies. The total amount of protein was visualized by Coomassie staining. **b.** Enzymatic assay performed with a range of different CCDC6-RET concentrations (7.5, 5 and 2.5 μM) incubated with increasing concentrations of ATP. Data represent the mean ± SEM of two experiments in duplicate (n = 4). Enzymatic activity (OD s^−1^ × 10^−3^). Catalytic efficiency constants (k_cat_/K_M_, fold difference) are depicted in the panel below. **c.** WB of samples from a time-course phosphorylation assay with using CCDC6-RET (1 μM, enzyme) in the presence of an inactive ∇CCDC6-RET-HRA mutant construct (3 μM, substrate surrogate); and c-Src (3D, 1 μM enzyme) in the presence of a c-Src KD K298M (3 μM, substrate), respectively; using the indicated antibodies. The total amount of protein was visualized by Coomassie staining. **d.** CCDC6-RET (5-6 μM), apo and autophosphorylated (30 min) samples were cross cross-linked with DSBU as indicated in material and methods. Samples were run on SDS-PAGE and stained with Coomassie, left panel. Intermolecular cross-linked peptides specific for the apo (0P, purple) and autophosphorylated (P, green) states are depicted schematically in color code, right panel. **e.** Intermolecular cross-linked peptides were filtered by distances between 12-31 Å with different 3D-structural models defining three different configurations: apo (0P, trans-inhibited, model 1), intermediate (0P, transition, model 2) and autophosphorylate (P, active, model 3) states. **f.** Proposed model for the mechanisms of CCDC6-RET autoactivation. CCDC6-RET adopts a face to face transinhibited configuration in the apo state characterized by cross-linked activation loops and αC helices facing each other. Upon nucleotide binding and catalytic domains reorientation (intermediate or transient state), CCDC6-RET undergoes *cis*-autophosphorylation (active, phosphorylated state).

Cross-linking MS is a technique that captures conformational changes in solution by mapping structural details of individual proteins and their complex assemblies based on the coordination of amino acid pairs that are positioned in close proximity to each other by using chemical crosslinkers (Iacobucci et al., 2018). These covalent connections then serve as basis for deriving distance constraints within the protein(s) under investigation. To provide experimental restraints for the 3D-reconstruction of the different functional states of a CCDC6-RET homodimer (apo and autophosphorylated), we performed in-solution cross-linking with the heterobifunctional cross-linker DSBU (disuccinimidyl dibutyric urea) (Iacobucci et al., 2018). Cross-linking was performed under apo and saturating ATP and MgCl_2_ (60 min) conditions (Fig. 8d). Mass spectrometry analysis of cross-linked residues in both apo and fully phosphorylated states (Fig. 8d) revealed several significant changes upon phosphorylation (Fig. 8d and supplemental table 4). The apo inactive state was defined by the presence of several intermolecular crosslinked peptides that connected the activation loop (K907-K907); the β2 of the G-loop (K737-K737) and the c-terminal part of the coiled-coil interface (S98-S98) of both kinases. These intermolecular connections were not observed in the phosphorylated form of the dimer, which was further defined by the presence of a crosslinked peptide between the final section of the coiled-coil interface and the β1 of the GRL (T100-K715, see Supplemental table 4). These results are compatible with a face-to-face arrangement of both RET kinase domains within the dimer in the apo (inactive) and transient states (Fig. 8e). In the presence of nucleotide and MgCl_2_, the catalytic domains swing apart (as indicated by the breaking up of the activation loop crosslinked peptide) adopting a 2-fold rotational symmetry compatible with a *cis* autophosphorylation mechanism. The reorientation of the catalytic domains is coordinated with the motion of the coiled-coil interface in a way that resemble a “chopstick” model, where a tighter base at the coiled-coil grasp the two catalytic domains together into a compacted transinhibited face-to-face dimer (Fig. 8e and f). In the active state, the contacts at the base of the coiled-coil interface are loosen, resulting consequently in a more relax and open interface at the base of the coiled-coil and a more proximal amino-terminal sections of the dimerization interface compatible with the motion (swinging) of the catalytic domains (Fig. 8e and f). Together the biochemical and cross-linking mass spectrometry data support a model for a face to face dimer compatible with a transinhibited state in the apo form. Upon nucleotide binding, autoactivation takes place via a *cis* mechanism driving CCDC6-RET autophosphorylation, (Fig. 8e and f).

## Discussion

In this work we uncover the molecular and structural determinants that control the autoactivation mechanism of a CCDC6-RET fusion product, a driver and therapeutic target in lung adenocarcinoma and papillary thyroid cancer (Kohno et al., 2012; Lipson et al., 2012; Wang et al., 2012). The RET/PTC1 (i.e. CCDC6-RET) oncogene was discovered more than 30 years ago by the lab of Giancarlo Vecchio (Fusco et al., 1987), and despite of a significant amount of published studies on the genetics and signalling of this oncogene and oncoprotein respectively, the molecular and structural bases for its (oncogenic) mechanism of action have remained largely elusive. Here, we biochemically dissected the autoactivation mechanism and found that CCDC6-RET is a highly active dimeric kinase in solution (Fig. 1). We found that the CCDC6-RET fusion product functions not only as a kinase with dual specificity for tyrosine and serine residues (Fig. 2); but also, strikingly as a dual ATP- and ADP-dependent kinase, able to bind and use both nucleotides as phosphoryl donors (Fig. 3). We also uncovered a catalytic dependency on activation loop Tyr 905 in the absence of the c-terminal segment (Fig. 5). These results were unexpected when compared with our previously published studies demonstrating that wild-type RET kinase or intracellular domains constructs lacking activation loop Tyr 905 (Y905F) did not compromise the catalytic activity in autophosphorylation experiments nor in enzymatic assays using substrate peptides (Knowles et al. 2006, Plaza-Menacho et al. 2014, Plaza-Menacho et al. 2016). Furthermore, our data suggest that the other RET activation loop tyrosine site, Tyr 900, is required for the adequate synthesis and folding of the protein in a process that appears to be dependent on the integrity of the whole catalytic core of the fusion and in particular of the c-terminal segment (Supplemental Fig. 3). Altogether, these findings make us to conclude that a yet unidentified element from the c-terminal segment makes key contacts with (phosphorylated) Tyr 900 at the activation loop in the apo state that are important for the stability and folding of the protein. Such detrimental impact caused by Tyr 900 mutation is strikingly rescued by the concomitant mutation of the other activation loop Tyr 905 autophosphorylation site. Interestingly, activation loop Tyr 900 appears already phosphorylated at time zero in the apo state, and upon CCDC6-RET autoactivation its phosphorylation levels correlate inversely with the phosphorylation state of activation loop Tyr 905, which undergoes fast and robust autophosphorylation in a time dependent manner, see Fig. 2. In this active setting, phosphorylation on c-terminal sites will release the c-tail to act as a scaffold or signalling platform to assemble signalling complexes. This is also supported also by the fact that a c-terminal deleted construct appears to be more efficient catalytically (Fig. 4). We hypothesize that the lack of availability of the c-terminal segment mimics its own engagement in the signalling assembly and transduction, where the activation loop Tyr 905 is catalytic required. This comes in line with our proposed mechanism of self-activation *in cis*-based on our biochemical (Fig. 8) and structural (Fig. 6 and 7) data. Furthermore, we generated a 3D-assembly of a CCDC6-RET homodimer combining an integrated structural biology and MS approach. We biochemically demonstrated that in accordance to our structural reconstruction, CCDC6-RET adopt a face-to-face (trans-inhibited) dimer in the apo-state, and upon nucleotide binding and conformational readjustment activation loop autophosphorylation is driven by a mechanism *in cis* (see Fig. 8 and 9). These data were unexpected in first place, as we anticipated that a constitutive (forced) protein kinase dimer would undergo direct intermolecular (i.e. *in trans*) autophosphorylation; but at the same time compatible with the domain arrangement seen in bacterial histidine kinases, where a catalytic domain situated to the left of the coiled-coil helix b (i.e. to the left of the four-helix bundle and 2-fold symmetry plane) undergoes *cis*-autophosphorylation (Bhate et al., 2015). Recent studies demonstrated that PKD autoinhibition in trans is compatible with and controlled by an activation loop autophosphorylation mechanisms in cis (as a monomer) (Reinhardt et al., 2023). In another relevant study, they show that PDK1 is autoinhibited by its PH domain intramolecularly (*in cis*) in the apo state and that upon PIP3 binding positive cooperativity promotes a switch-like activation of PDK1 driving intermolecular PDK1 autophosphorylation by the assembly of a face-to-face dimer (Levina et al., 2022). Our results are in line with these recent studies that highlight altogether the existence of complex regulatory elements during the autoactivation mechanism, which are controlled by intra- and inter-molecular components that appears to be rather specific and private for each kinase (Reinhardt and Leonard, 2023). These components can interconvert from case to case depending on the accessory domains to the catalytic core and the signalling inputs that control that particular kinase. From a biological point of view, the dual ATP and ADP dependency seen by the CCDC6-RET fusion product would result in a gain of functionality by being able to perform two catalytic cycles from one single ATP molecule; transferring both the γ- and β- phosphate groups of the initial ATP and its immediate product ADP, respectively, onto a susbtrate and/or different susbtrates surrogates. In the cell, the ATP concentration is maintained in the range of 1 to 10 mM, with a normal ratio of ATP/ADP of approximately 1000. In order to overcome such concentration barrier, the local concentration of the ADP source has to be coupled with the formation of CCDC6-RET protein condensates and lipid phase separation therefore localizing and focalizing the ADP dependent signal (Shelby et al., 2023; Wang et al., 2023). In this line, we are currently undertaking substrate identification in cells (Knight et al., 2012) using a whole-cell lysate kinase assay coupled with LC/MS-MS under saturating concentrations of both nucleotides ATP and ADP (work in preparation). Our preliminary results indicate that CCDC6-RET exert a different pattern of phosphorylated susbtrates in cells; also showing a higher activity (i.e. higher number of phosphorylated susbtrates identified) when ADP was used to trigger the phosphorylation reaction as compared to ATP. The incremental number of phosphorylated peptide-susbtrates is explained however not by an increase in the phospho-tyrosine activity of CCDC6-RET, but by a large increment in the appearance of Ser-containing phosphorylated susbtrates. These data suggest that ADP-dependent activity by the CCDC6-RET fusion product can be coupled to a change in susbtrate specificity towards Ser rather than Tyr; in other words, ADP-dependency appears to be associated with a change in the dual specificity toward Ser phospho-sites. The dual ATP- and ADP-dependency seen by CCDC6-RET is a striking mechanistic feature, not previously described by any other protein kinase as far as we are aware, as usually ADP-dependent kinases are very restrictive to their ligands being unable to use tri-phosphorylated nucleotides such as ATP (Guixe and Merino, 2009; Ito et al., 2001; Richter et al., 2016). However, it has been shown that they can bind ATP by competition kinetic experiments (Guixe and Merino, 2009). This dual ATP/ADP dependency could have important implications for how cells maintain signaling functions under stressful conditions such as low oxygen or ischemia, which are common in diseases like cancer and atherosclerosis.

There is a striking lack of homology and fold-conservation between RET (and human protein kinases in general) and human ADP-dependent kinases, taking for example the ADP-dependent glucokinase (ADPGK) as a main paradigm (Richter et al., 2016). ADP-dependent kinases are classified as members of the ribokinase superfamily (Guixe and Merino, 2009; Ito et al., 2001). The ribokinase-like fold is basically composed of an eight-stranded β-sheet surrounded by eight α-helices, three on one side and five on the other which constitutes a single domain (the large domain). This superfamily initially included ATP-dependent kinases of adenosine, fructose, tagatose-6-P, fructose-6-P, and fructose-1-P among others, besides ribokinase, the canonical enzyme (Bork et al., 1993). Later, the ribokinase superfamily also included enzymes that can transfer the γ-phosphate of ATP to some vitamins involved in B6 synthesis, such as pyridoxal kinase (Li et al., 2002). This superfamily can be subdivided into three major groups: the ATP-dependent vitamin kinases, the ATP-dependent sugar kinases, and the ADP-dependent sugar kinases. The main structural difference between these groups is related to the presence of a second (small) domain composed by a β-sheet, with some α-helical insertions in the case of the ADP-dependent kinases and ATP-dependent kinases; while the vitamin kinase enzymes present only the αβα ribokinase-like fold (the large domain). Leucine-zipper motifs are one of the main structural motifs known to interact with DNA elements Note that the coiled-coil interface of CCDC6-RET resembles highly of a leucine-zipper motif characterized by a Leu in position d of the heptad repeat; which in the case of the RET fusion is formed by Leu 59, Leu 66, Leu 73, Tyr 80, Leu 87, Leu 94 and Ile 101. The connection of CCDC6-RET with the DNA-damage response pathway, its interaction with DNA-elements and protein kinases and other effectors implicated in this fundamental process are yet to be elucidated. Interestingly, the RET S909 autophosphorylation site is within an ATR/ATM (Plaza-Menacho et al., 2016) consensus site (pS/TQG), being potentially an intermolecular regulatory element. Contrary to the case of the wild-type RET (Plaza-Menacho et al., 2016), this residue when mutated appears to have a profound effect on the catalytic activity of the RET fusion product towards a substrate despite lacking any effect on autophosphorylation (unpublished data). This is in fact a clear evidence that autophosphorylation and the phosphorylation of an exogenous susbtrate (e.g. a peptide, or an intact susbtrate surrogate) are not governed by the same mechanistic determinants and despite being both used as readouts for catalytic activity, they should be treated as independent mechanisms or processes (Plaza-Menacho et al., 2016; Plaza-Menacho et al., 2014b; Reinhardt and Leonard, 2023). Our biochemical data allowed us to discriminate between *cis*- and *trans*-autophosphorylation components by means of using active protein kinases that can (i.e. c-Src) or cannot (i.e. CCDC6-RET) phosphorylate intermolecularly an inactive susbtrate surrogate that mimics its own kinase domain (Fig. 8c). The crosslinking mass spectrometry data filtered under our 3D-reconstructed structural models recapitulate the different conformational and functional states in the autoactivation process of CCDC6-RET (Fig. 8d-f): i) the apo state characterized by a face to face arrangement of both catalytic domain captured by a critical intermolecular crosslinked peptide between both activation loops, and a closer interface at the base of the coiled-coil; ii) transition state upon nucleotide binding where the catalytic domains start to swing away (swap) from each other and the contacts at the base of the coiled-coil are less tighter, and iii) active autoactivation state, defined by catalytic domain swap and cis-auto-phosphorylation and opening of the coiled-coil base (Fig. 8f). Despite the resolution limitation of our EM data (negative staining), our integrative approach provides altogether a feasible tridimensional representation of the twisted arrangement of the catalytic domains from the coiled-coil interface, in line with our proposed model of cis-phosphorylation. We are currently trying to obtain higher resolution information by cryo-EM studies that will allow us to solve high-resolution structures of the different functional/conformational steps/stages to fully support our mechanistic studies. All in all, our work presented uncovers for the first time the molecular and structural determinants that controls CCDC6-RET function and provides a solid framework for the structure-function analyses of other RET fusion products.

## Author contributions

Conception of the study, experimental design, supervision of the study, funding acquisition and preparation of first draft (I.P.-M.), manuscript writing, data analysis and figures preparation (I.P.-M., A.M.-H.), mass spectrometry including data analysis (J.S.-W., J.M., E.Z and M.I.), SAXS studies including data analysis and figure preparation (I.M.), experimental work and data acquisition (A.M.-H., J.C., I.P.-M.).

## Acknowledgements

We are grateful to: Jasminka Boskovic and Johanne Le Cop from the Electron Microscopy Unit (CNIO), Fernando García from the Proteomics Unit (CNIO), the Genomics Unit (CNIO), former members of the Protein Phosphorylation and Cancer Group for their contribution at the very early stages of the project (Pablo Soriano, Nicolás Cuesta y Julio Martínez-Torres). We thank Nabil Djouder for helpful comments and advice on the manuscript. The authors would like to thank Diamond Light Source for beamtime (beamline B21, proposal mx30297) and their staff for assistance during data collection. We thank the Centro Nacional de Investigaciones Oncológicas (CNIO), which is supported by the Instituto de Salud Carlos III and recognized as a “Severo Ochoa” Centre of Excellence (ref. CEX2019-000891-S, awarded by MCIN/AEI/ 10.13039/5 01100011033) for core funding and supporting this study. This work was further supported by projects from the Spanish Ministerio de Ciencia e Innovación (MCIN), BFU2017-86710-R funded by MCIN/AEI /10.13039/501100011033 and ERDF “A way of making Europe”, PID2020-117580RB-I00 funded by MCIN/ AEI /10.13039/501100011033, RYC-2016-1938 funded by MCIN/AEI /10.13039/501100011033 and ESF “Investing in your future”, and a FP7-PEOPLE-2013-COFUND - Marie-Curie Action: “Co-funding of regional, National and International Programmes” International grant (number 608765) to I.P.-M.

