## Supplemental material and figures for "The oncogenic CCDC6-RET fusion product is a dual ATP and ADP-dependent kinase that functions via *cis*-phosphorylation"

### 1. Supplemental figures

Supplemental figure 1. Functional characterization of CCDC6-RET phospho-site mutants

Supplemental figure 2. Response to different nucleotides (ATP, ADP and AMP) by CCDC6-RET.

Supplemental figure 3. Purification and characterization of a CCDC6-RET Y900F mutant

Supplemental figure 4. Structural inspection of the in silico assembled CCDC6-RET dimer and generation of CCDC6-RET dimers using AlphaFold multimer models.

Supplemental figure 5. EM analyses of CCDC6-RET.

Supplemental figure 6. SAXs analyses of a CCDC6-RET ΔCT construct.

### 2. Supplemental tables

Supplemental table1. Phospho-proteomic identification of CCDC6-RET autophosphorylation sites by LC/MS-MS.

Supplemental table2. Phospho-proteomic characterization of ADP-dependent CCDC6-RET activity by LC/MS-MS.

Supplemental table 3. SAXs data collection and structural and mass parameter of a CCDC6-RET ΔCT construct.

Supplemental table 4. Intermolecular crosslinked peptides from the XL-MS analyses for CCDC6-RET apo and phosphorylated states.

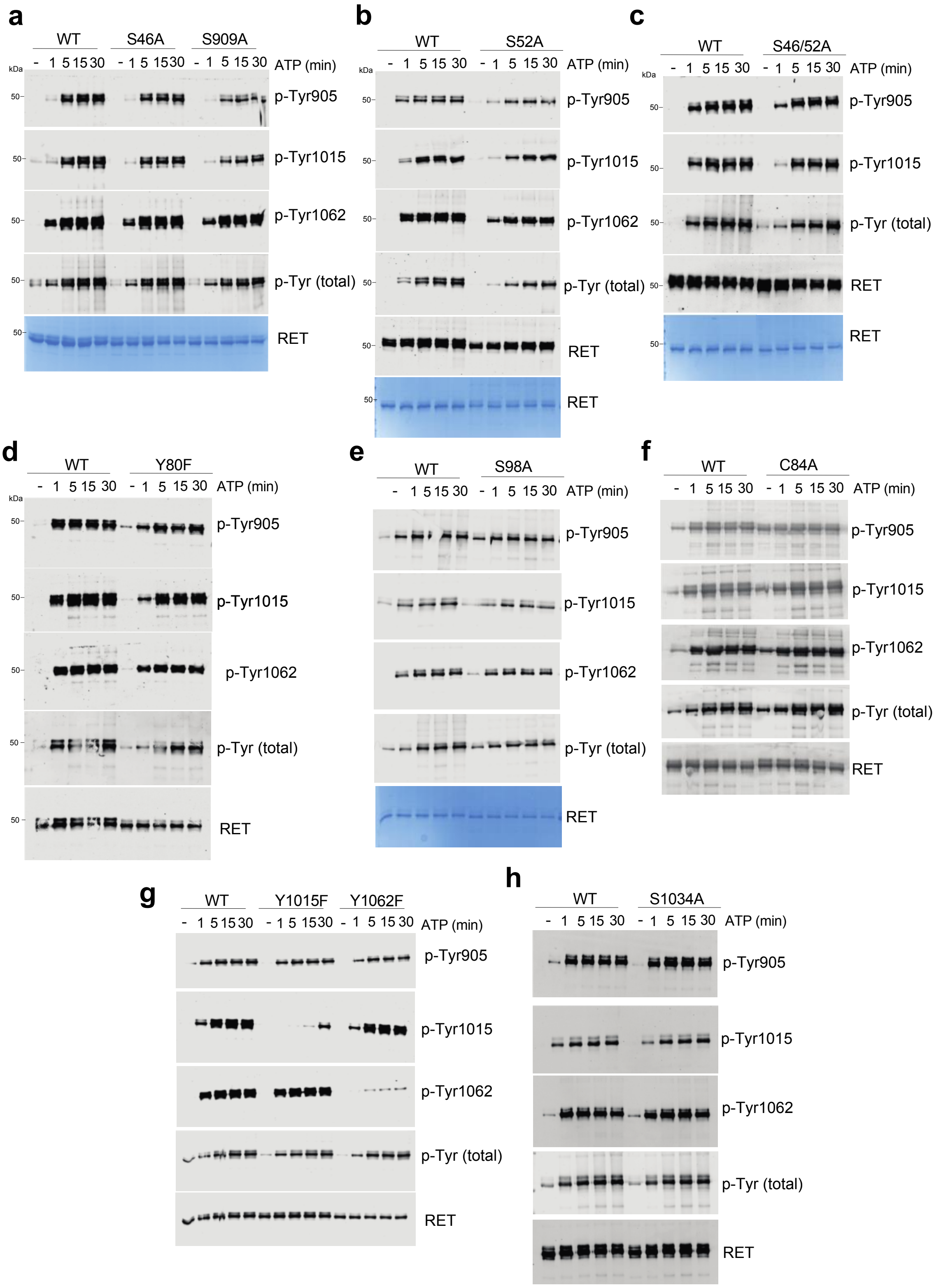


Supplemental Figure 1. Functional characterization of CCDC6-RET phospho-site mutants.

WBs analyses of time-course (0-30 min) autophosphorylation experiments of recombinant CCDC6-RET WT and the following phospho-site mutants: S46A and S909A (a), S52A (b), S46/52A (c), Y80F (d), S98A (e), C84A (f), Y1015F and Y1062F (g) and S1034 (h) using the indicated antibodies. Total amount of protein was visualized by Coomassie staining or by using a total RET antibody.

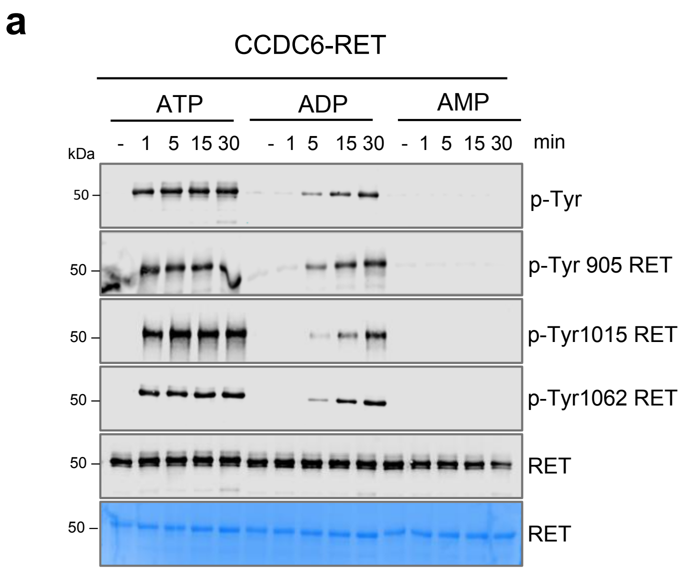


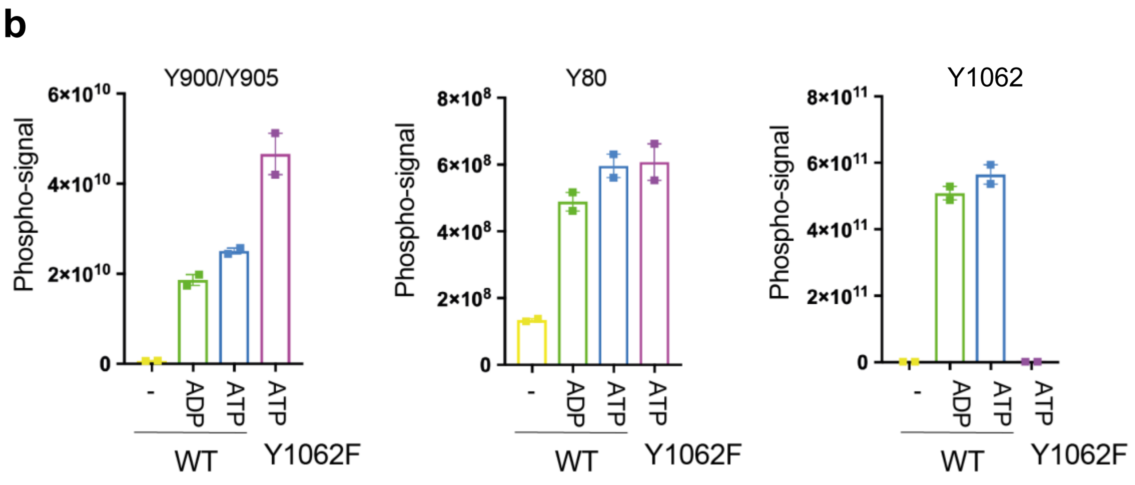


Supplemental figure 2. Response to different nucleotides (ATP, ADP and AMP) by CCDC6-RET.

a. WBs analyses of a time-course (0-30 min) autophosphorylation experiment of recombinant CCDC6-RET (1-2 μM) using different nucleotides (1 mM) AMP, ADP and ATP in presence of 2 mM of $MgCl_2$.

b. LQ/MS-MS analyses of CCDC6-RET WT (1-2 μM) stimulated with ATP and ADP (1 mM) in presence of $MgCl_2$ (2 mM) at 0 (unstimulated) and 30 min. A CCDC6-RET Y1062F mutant in the presence of ATP (1 mM) in and $MgCl_2$ (2 mM) was also evaluated in the same experimental conditions. Representative phospho-peptides are depicted.

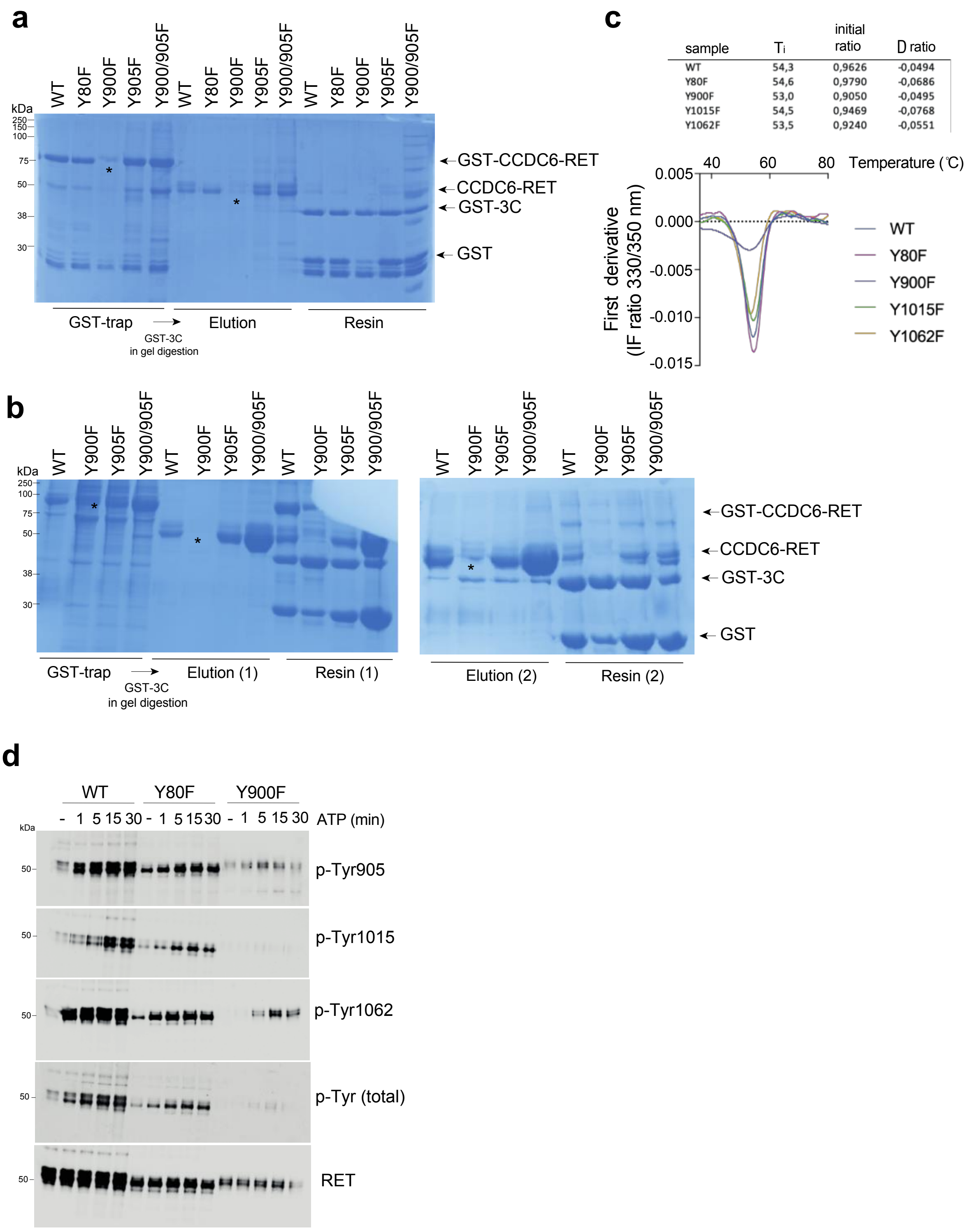


| sample | $T_i$ | initial ratio | D ratio |
|---|---|---|---|
| WT | 54,3 | 0,9626 | -0,0494 |
| Y80F | 54,6 | 0,9790 | -0,0686 |
| Y900F | 53,0 | 0,9050 | -0,0495 |
| Y1015F | 54,5 | 0,9469 | -0,0768 |
| Y1062F | 53,5 | 0,9240 | -0,0551 |

Supplemental figure 3. Purification and characterization of a CCDC6-RET Y900F mutant.

**a-b.** Samples of recombinant CCDC6-RET (WT and indicated mutants) from the different protein purification steps: recombinant tagged proteins bound to Glutathione conjugated resin, eluted samples after 3C-protease treatment and resin after elution. Protein were visualized by SDS-PAGE and Coomassie staining.

**c**. DSF analyses by direct IF (tycho nanotemper) of eluted (soluble) samples from a providing thermal shifts changes ($T_i$), initial ratio and $\Delta$ratio values, respectively.

**d**. WB analyses of a time-course (0-30 min) autophosphorylation experiment using CCDC6-RET (1-2 $\mu$M) WT, Y80F and Y900F in presence of ATP (1 mM) and 2 mM of $MgCl_2$, using the indicated antibodies.

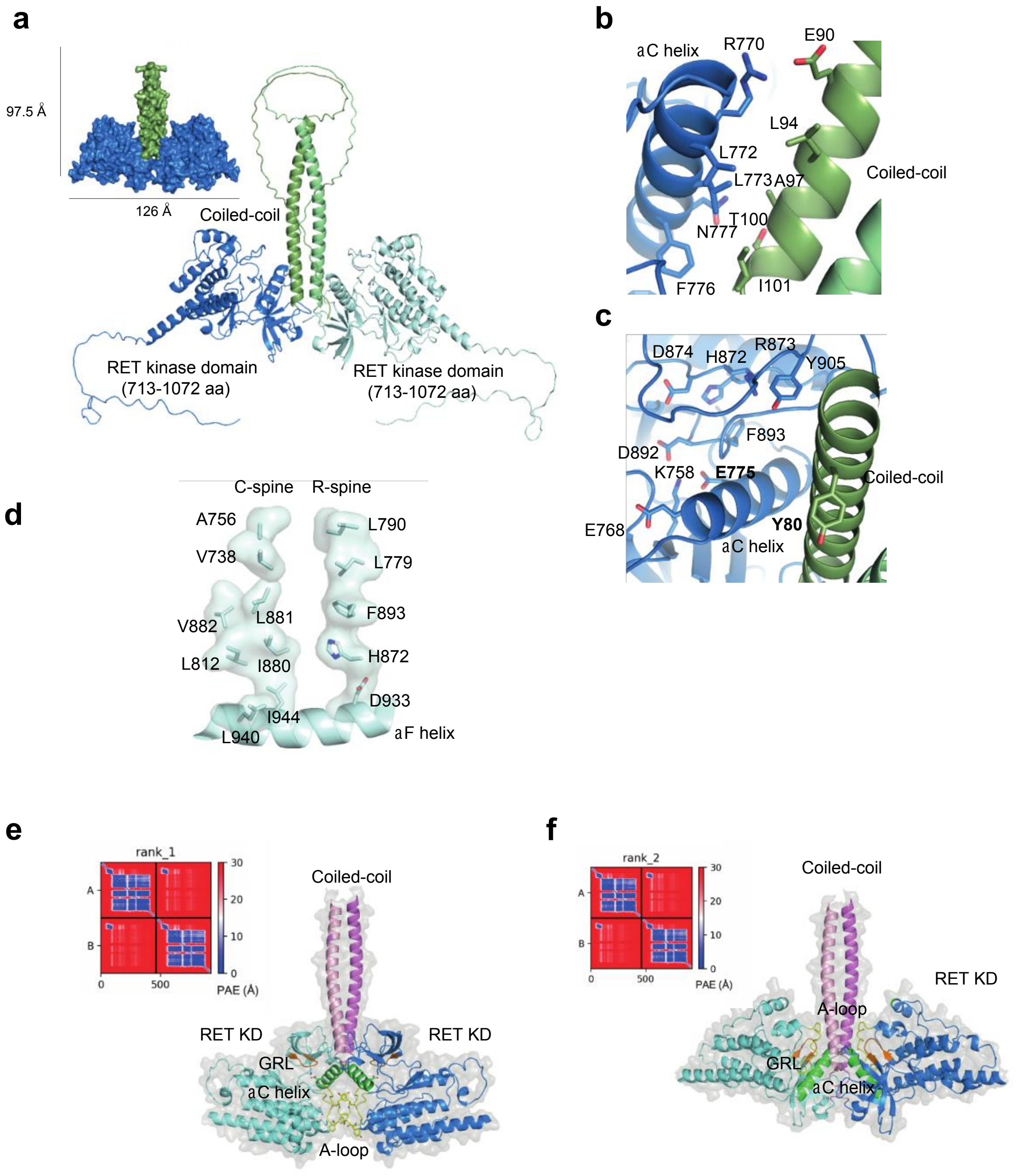


Supplemental figure 4. Structural inspection of *in silico* assembled CCDC6-RET homodimers.

**a**. *In silico* reconstruction of a CCDC6-RET homodimer from a high-confidence 3D-predictive model Alpha-Fold (AF id: J7M8C2), see figure 6. Indicated particle size (Å) visualized in surface representation.

**b**. Cartoon representation with key side chain residues depicted in sticks from the hydrophobic patch between the PIF-like pocket at the top of the αC helix and the coiled-coil.

**c**. Cartoon representation with key side chain residues depicted in sticks from the active site HRD (872-874), DFG (892-894) and catalytic salt-bridge (K758-E775) are depicted, aa numbering of RET native sequence.

**d**. View of the linear architecture of the catalytic (C) and regulatory (R) spines of CCDC6-RET featuring an active DFG-in configuration. Side chains of aa composites are depicted, numbering of aa follow the RET native sequence.
**e-f**. Per-residue confident score (pLDDT) of the top two generated models by AlphaFold multimer, ranked 1 and 2 and predicted aligned error (PAE) heat map of both top ranked models. The predicted error between all pairs of residues is shown in a gradient blue-red scale for each monomer molecule. Cartoon (color coded, pink coiled-coil, blue kinase core, green aC helix, orange G-loop, and yellow A-loop, monomer A softer color code) and surface (grey, transparent) representation of predicted models without amino- and carboxy-terminal IDRs sequences.

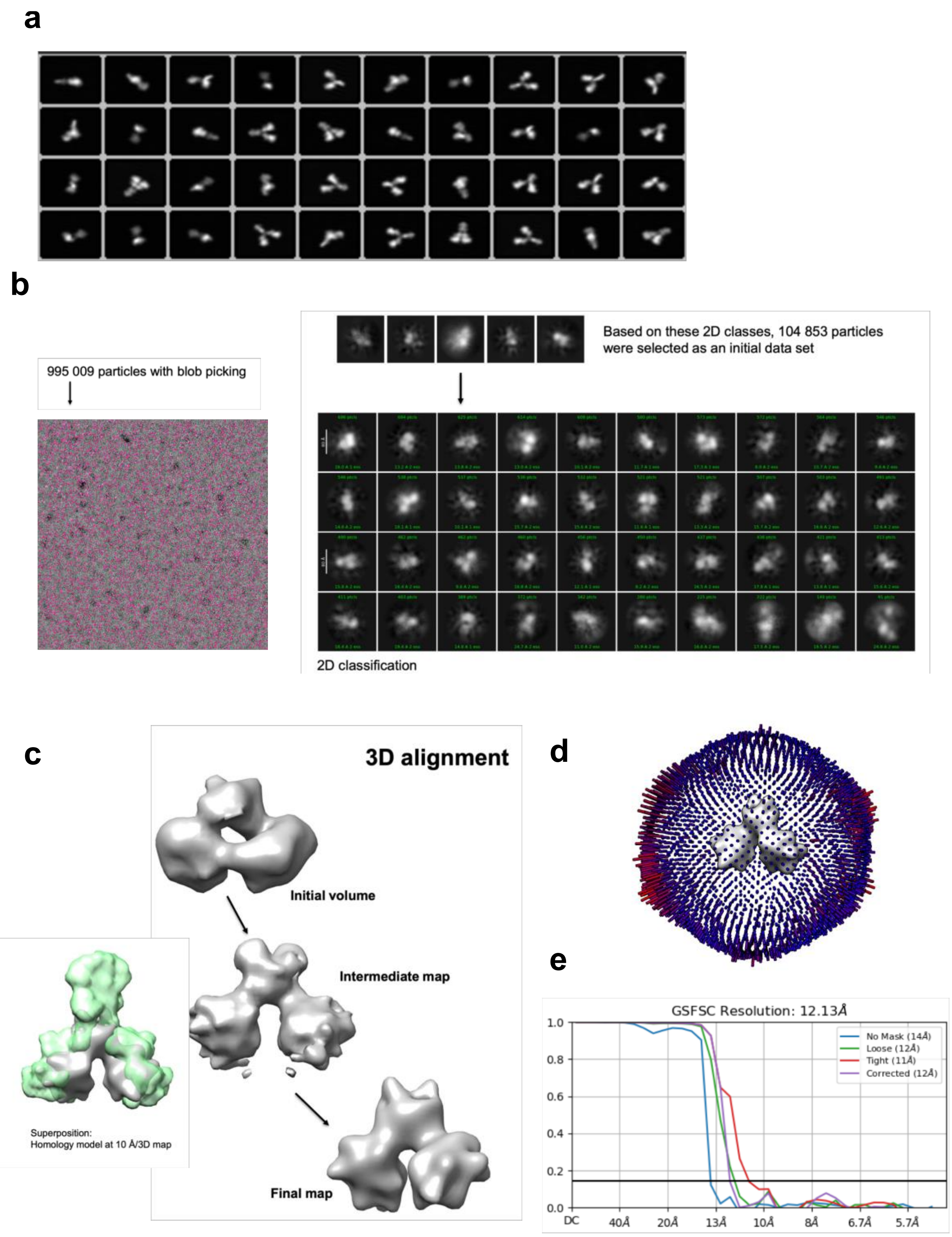


Supplemental figure 5. EM analyses of CCDC6-RET.

a. Retro-projections of a homodimer CCDC6-RET 3D-model adopting an inverted Y shape.

b. Particle acquisition and data processing with cryoSPARK. NS micrograph particle selection (995.009) with blob picking, an initial 2D classification was made, and based on this, 108.853 particles were selected to build an initial dataset with the final 2D-classification depicted.

**c**. We generated an initial 3D-map for the dataset, which we refined further obtaining intermediate and final maps. The reliability of the map is attested by the good agreement between classes and reprojections (see a), the even distribution of Euler angles (d)

**d**. Euler angles distribution from c.

**e**. Resolution estimate by Fourier shell correlation.

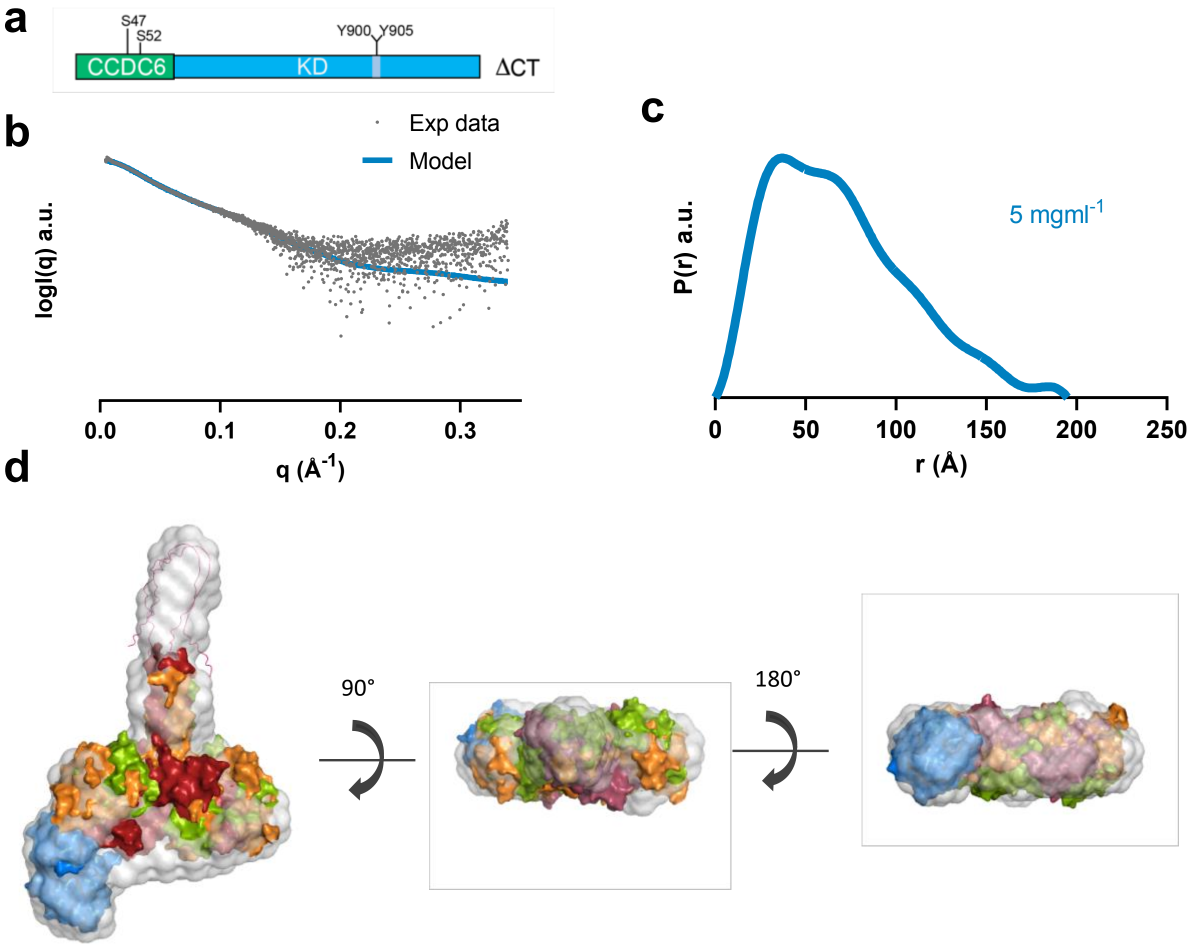


Supplemental figure 6. SAXS analyses of a CCDC6-RET ΔCT construct.

a. Schematic diagram of a CCDC6-RET ΔCT construct used in our studies depicting functional domains with the indicated phospho-sites and aa number.

b. Theoretical and empiric scattering curves of a CCDC6-RET ΔCT protein (5 $mgml^{-1}$). The fitting of the theoretical curves derived from CCDC6-RET ΔCT model against the experimental curves gave a $\chi^2$ value of 2.2

c. Pair distribution function from b.

d. Unbiased ab initio model superimposed with the three different 3D-*in silico* reconstructed models (see figure 6 and supplemental figure 4). Note, the models (manual: orange, model 2: red, model 5: green) do not include the amino nor the carboxy-terminal IDRs sequences. Amino terminal region of CCDC6-RET (aa 1-52) was manually modelled inside the volume solution.

| Phosphopeptide Sequence (charge) | Protein name | Phosphosite |
|---|---|---|
| ADSASESDTDGAGGNSSSSAAMQSSCSSTSGGGGGGGGGGGGGK (+3) | CCDC6 | S4 |
| MADSASESDTDGAGGNSSSSAAMQSSCSSTSGGGGGGGGGGGGGK (+3) | CCDC6 | S6 |
| SGGIVISPFR (+2)<br>SGGIVISPFRLEELTNR (+2) (+3) | CCDC6 | S46 |
| SGGIVISPFR (+2)<br>SGGIVISPFRLEELTNR (+2) (+3)<br>SGGIVISPFRLEELTNRLASLQQENK (+4) | CCDC6 | S52 |
| LASLQQENK (+2)<br>LASLQQENKVLK (+2) | CCDC6 | S65 |
| LASLQQENKVLKIELETYK (+3) (+4)<br>VLKIELETYK (+2) (+3)<br>VLKIELETYKLK (+2) (+3) (+4)<br>IELETYK (+2)<br>IELETYKLK (+2) (+3) | CCDC6 | Y80 |
| KASVTIEDPK (+2)<br>KASVTIEDPKWEFPR (+2) (+3) (+4)<br>ASVTIEDPKWEFPRK (+3) | CCDC6 | S98 |
| QVNHPHVIKLYGACSQDGPLLLIVEYAK (+4)<br>LYGACSQDGPLLLIVEYAK (+2) (+3) | RET | Y791 |
| QVNHPHVIKLYGACSQDGPLLLIVEYAK (+4)<br>LYGACSQDGPLLLIVEYAK (+2) (+3) | RET | Y806 |
| YGSLRGFLR (+2) | RET | Y809 |
| ESRKVGPGYPDER (+3)<br>KVGPGYPDER (+2) (+3)<br>VGPGYPDER (+2) | RET | Y826 |
| ISDFGLSRDVYEEDSYVKR (+3) | RET | S896 + Y900 |
| ISDFGLSRDVYEEDSYVKR (+3) | RET | S896 + Y900 + Y905 |
| ISDFGLSRDVYEEDSYVK (+2) (+3)<br>ISDFGLSRDVYEEDSYVKR (+2) (+3) (+4)<br>DVYEEDSYVK (+2)<br>DVYEEDSYVKR (+2) (+3) | RET | Y900 |
| ISDFGLSRDVYEEDSYVKR (+2) (+3) (+4)<br>DVYEEDSYVKRSQGR (+2) (+3) | RET | Y900 + S904 |
| ISDFGLSRDVYEEDSYVKR (+3)<br>DVYEEDSYVKR (+2) | RET | Y900 + S904 +Y905 |
| MKISDFGLSRDVYEEDSYVK (+3)<br>ISDFGLSRDVYEEDSYVK (+2) (+3)<br>DVYEEDSYVK (+2)<br>DVYEEDSYVKR (+2) (+3) | RET | Y900 + Y905 |
| DVYEEDSYVKRSQGR (+3) | RET | S904 |
| ISDFGLSRDVYEEDSYVK (+2) (+3)<br>ISDFGLSRDVYEEDSYVKR (+2) (+3) (+4)<br>DVYEEDSYVK (+2)<br>DVYEEDSYVKR (+2) (+3) | RET | Y905 |
| SQGRIPVK (+2) | RET | S909 |
| TGHRMERPDNCSEEMYR (+3) (+4)<br>MERPDNCSEEMYR (+2) (+3)<br>MERPDNCSEEMYRLMLQCWK (+3) (+4)<br>PDNCSEEMYR (+2) | RET | Y981 |
| RRDYLDLAASTPSDSLIYDDGLSEEETPLVDCNNAPLPR (+3) (+4)<br>RDYLDLAASTPSDSLIYDDGLSEEETPLVDCNNAPLPR (+3)<br>DYLDLAASTPSDSLIYDDGLSEEETPLVDCNNAPLPR (+3) (+4) | RET | Y1015 |
| DYLDLAASTPSDSLIYDDGLSEEETPLVDCNNAPLPR (+3)<br>DYLDLAASTPSDSLIYDDGLSEEETPLVDCNNAPLPR (+4) | RET | Y1015 +S1021 |
| RRDYLDLAASTPSDSLIYDDGLSEEETPLVDCNNAPLPR (+4)<br>RDYLDLAASTPSDSLIYDDGLSEEETPLVDCNNAPLPR (+3) (+4) | RET | Y1015 +Y1029 |
| RDYLDLAASTPSDSLIYDDGLSEEETPLVDCNNAPLPR (+3) (+4) | RET | Y1015 +S1034 |
| RRDYLDLAASTPSDSLIYDDGLSEEETPLVDCNNAPLPR (+3) (+4)<br>DYLDLAASTPSDSLIYDDGLSEEETPLVDCNNAPLPR (+3) (+4) | RET | Y1029 |
| RRDYLDLAASTPSDSLIYDDGLSEEETPLVDCNNAPLPR (+4)<br>RDYLDLAASTPSDSLIYDDGLSEEETPLVDCNNAPLPR (+4) | RET | Y1029 + S1034 |
| RRDYLDLAASTPSDSLIYDDGLSEEETPLVDCNNAPLPR (+3) (+4)<br>RDYLDLAASTPSDSLIYDDGLSEEETPLVDCNNAPLPR (+3) (+4)<br>DYLDLAASTPSDSLIYDDGLSEEETPLVDCNNAPLPR (+3) (+4) | RET | S1034 |
| ALPSTWIENKLYGR (+2) (+3) | RET | Y1062 |

Supplemental table 1. Phospho-sites identification by LC/MS-MS during the CCDC6-RET autophosphorylation reaction *in vitro* in presence of ATP (1 mM) and $MgCl_2$ (2 mM), see Fig. 2.

| | Protein name | Phosphosite |
|---|---|---|
| SGGIVI**S**PFRLEELTNR (+2) (+3)<br>SGGIVI**S**PFRLEELTNRLASLQQENK (+3) (+4) | CCDC6 | S52 |
| VLKIELET**Y**KLK (+2) (+3)<br>IELET**Y**KLK (+2) (+3)<br>IELET**Y**KLKCK (+2) (+3) (+4) | CCDC6 | Y80 |
| QVNHPHVIKL**Y**GACSQDGPLLLIVEYAK (+3) (+4) | RET | Y791 |
| **Y**GSLRGFLR (+2) (+3)<br>YGSLRGFLRESR (+3) | RET | Y809 |
| LYGACSQDGPLLLIVEYAKYG**S**LRGFLR (+3) (+4) | RET | S811 |
| ESRKVGPG**Y**PDER (+2) (+3)<br>KVGPG**Y**PDER (+2) (+3)<br>VGPG**Y**PDER (+2)<br>KVGPG**Y**PDERALTMGDLISFAWQISQGMQYLAEMK (+3) (+4) | RET | Y826 |
| ISDFGL**S**RDVYEEDSYVKR (+2) (+3) (+4) | RET | S896 |
| ISDFGL**S**RDV**Y**EEDSYVKR (+3) | RET | S896 + Y900 |
| ISDFGL**S**RDV**Y**EEDS**Y**VKR (+3) | RET | S896 + Y900 + Y905 |
| ISDFGLSRDV**Y**EEDSYVK (+2) (+3)<br>ISDFGLSRDV**Y**EEDSYVKR (+2) (+3) (+4)<br>DV**Y**EEDSYVKR (+2) (+3)<br>MKISDFGLSRDV**Y**EEDSYVK (+3) | RET | Y900 |
| ISDFGLSRDV**Y**EED**S**YVK (+3)<br>DV**Y**EED**S**YVKR (+2) (+3)<br>ISDFGLSRDV**Y**EED**S**YVKR (+2) (+3) (+4) | RET | Y900 +S904 |
| DV**Y**EEDS**Y**VKR (+2) (+3)<br>DV**Y**EEDS**Y**VKRSQGR (+3)<br>ISDFGLSRDV**Y**EEDS**Y**VKR (+2) (+3) (+4)<br>ISDFGLSRDV**Y**EEDS**Y**VK (+2) (+3) | RET | Y900 + Y905 |
| DVYEED**S**YVKRSQGR (+3)<br>ISDFGLSRDVYEED**S**YVKR (+2) (+3) (+4) | RET | S904 |
| ISDFGLSRDVYEED**SY**VKR (+3) | RET | S904 + Y905 |
| ISDFGLSRDVYEEDS**Y**VK (+2) (+3)<br>ISDFGLSRDVYEEDS**Y**VKR (+2) (+3) (+4)<br>DVYEEDS**Y**VKR (+2) (+3) | RET | Y905 |
| MERPDNCSEEM**Y**R (+2) (+3)<br>MERPDNCSEEM**Y**RLMLQCWK (+3) (+4) | RET | Y981 |
| RRD**Y**LDLAASTPSDSLIYDDGLSEEETPLVDCNNAPLPR (+3) (+4)<br>RD**Y**LDLAASTPSDSLIYDDGLSEEETPLVDCNNAPLPR (+3) (+4)<br>D**Y**LDLAASTPSDSLIYDDGLSEEETPLVDCNNAPLPR (+3) (+4) | RET | Y1015 |
| D**Y**LDLAA**S**TPSDSLIYDDGLSEEETPLVDCNNAPLPR (+3) (+4)<br>RRD**Y**LDLAA**S**TPSDSLIYDDGLSEEETPLVDCNNAPLPR (+4) | RET | Y1015 + S1021 |
| RRD**Y**LDLAAS**T**PSDSLIYDDGLSEEETPLVDCNNAPLPR (+4) | RET | Y1015 + T1022 |
| RD**Y**LDLAASTP**S**DSLIYDDGLSEEETPLVDCNNAPLPR (+4) | RET | Y1015 + S1024 |
| RRD**Y**LDLAASTPSD**S**LIYDDGLSEEETPLVDCNNAPLPR (+4) | RET | Y1015 + S1026 |
| RRD**Y**LDLAASTPSDSLI**Y**DDGLSEEETPLVDCNNAPLPR (+4)<br>RD**Y**LDLAASTPSDSLI**Y**DDGLSEEETPLVDCNNAPLPR (+3) (+4) | RET | Y1015 + S1029 |
| RD**Y**LDLAASTPSDSLIYDDGLEEETPLVDCNNAPLPR (+3) (+4)<br>RRD**Y**LDLAASTPSDSLIYDDGL**S**EEETPLVDCNNAPLPR (+4) | RET | Y1015 + S1034 |
| RRD**Y**LDLAASTPSDSLIYDDGLSEEE**T**PLVDCNNAPLPR (+4) | RET | Y1015 + T1038 |
| RD**Y**LDLAA**ST**PSDSLIYDDGLSEEETPLVDCNNAPLPR (+3) (+4) | RET | Y1015 + S1021+ T1022 |
| RD**Y**LDLAA**S**TPSDSLI**Y**DDGLSEEETPLVDCNNAPLPR (+3) (+4) | RET | Y1015 + S1021+ Y1029 |
| RD**Y**LDLAAS**T**P**S**DSLIYDDGLSEEETPLVDCNNAPLPR (+4) | RET | Y1015 + T1022 + S1024 |
| RD**Y**LDLAAS**T**PSDSLI**Y**DDGLSEEETPLVDCNNAPLPR (+3) (+4) | RET | Y1015 + T1022 + Y1029 |
| RD**Y**LDLAASTP**S**D**S**LIYDDGLSEEETPLVDCNNAPLPR (+4) | RET | Y1015 + S1024 + S1026 |
| RD**Y**LDLAASTPSDSLI**Y**DDGL**S**EEETPLVDCNNAPLPR (+3) (+4) | RET | Y1015 + Y1029 + S1034 |
| RD**Y**LDLAASTPSDSLIYDDGL**S**EEE**T**PLVDCNNAPLPR (+4) | RET | Y1015 + S1034 + T1038 |
| RRDYLDLAAS**T**PSDSLIYDDGL**S**EEETPLVDCNNAPLPR (+4) | RET | T1022 + S1034 |
| RDYLDLAASTPSDSLI**Y**DDGL**S**EEETPLVDCNNAPLPR (+3) (+4)<br>RRDYLDLAASTPSDSLI**Y**DDGL**S**EEETPLVDCNNAPLPR (+4) | RET | S1024 + S1034 |
| RRDYLDLAASTPSD**S**LIYDDGL**S**EEETPLVDCNNAPLPR (+4) | RET | S1026 + S1034 |
| RRDYLDLAASTPSDSLI**Y**DDGLSEEETPLVDCNNAPLPR (+3) (+4) | RET | Y1029 |
| RDYLDLAASTPSDSLI**Y**DDGL**S**EEETPLVDCNNAPLPR (+3) (+4)<br>RRDYLDLAASTPSDSLI**Y**DDGL**S**EEETPLVDCNNAPLPR (+4) | RET | Y1029 + S1034 |
| RRDYLDLAASTPSDSLIYDDGL**S**EEETPLVDCNNAPLPR (+3) (+4)<br>RDYLDLAASTPSDSLIYDDGL**S**EEETPLVDCNNAPLPR (+3) (+4) | RET | S1034 |
| ALPSTWIENKL**Y**GR (+2) (+3) (+4)<br>ALPSTWIENKL**Y**GRISHAFTR (+3) (+4)<br>L**Y**GRISHAFTRF (+2) (+3) (+4) | RET | Y1062 |
| ALPSTWIENKLYGRI**S**HAFTR (+3) (+4) | RET | S1066 |
| ISHAF**T**R (+2)<br>ISHAF**T**RF (+2) (+3) | RET | T1070 |

Supplemental table 2. Phospho-sites identification by LC/MS-MS during the CCDC6-RET autophosphorylation reaction *in vitro* in presence of ADP (1 mM) and $MgCl_2$ (2 mM), see Fig. 3.

| | |
|---|---|
| Data collection parameters | |
| Instrument | Diamond Light Source beamline B21 (HarwellCampus, UK) |
| Wavelength (Å) | 1 |
| q-range ($Å^{-1}$) | 0.0032-0.34 |
| Exposure time (s) and final number of sample frames | 3, 600 |
| Concentration (mg $ml^{-1}$) | 5 |
| Temperature (K) | 293 |
| Structural parameters | |
| Protein | CCDC6-RET ∆CT |
| Rg (Å) (from Guinier) | 55.07 ± 0.18 |
| Rg (Å) (from P(r)) | 55.39 ± 0.2 |
| Dmax (Å) | 195 |
| Porod volume, $V_p$ ($Å^3$) | 212900 |
| Molecular mass (MM) determination | |
| Calculated MM (kDa) from sequence | 43,085 monomer / 86,07 dimer |
| From SAS, concentration independent method (MoW) | 125,23 dimer |
| From SAS-independent measure (SEC-MALS, kDa) | 85.0 ± 0.1 |
| Software employed | |
| Data processing | Scåtter/PRIMUS/ GNOM |
| Ab initio analysis / Averaging | DAMMIF, DAMMIN/DAMAVER |
| Computation of model intensities | FoXS |
| 3D graphics representations | PyMOL |

Supplemental table 3. SAXs data collection and structural and mass parameters for a CCDC6-RET ∆CT construct

| Crosslinked residues | Crosslinked peptides | Abundance in 0P state | Model #1 | Model #2 | Model #3 |
|---|---|---|---|---|---|
| K821— K821 | KVGPGYPDER—KVGPGYPDER (3)(4) | 9 | - | - | - |
| K907— K907 | DVYEEDSYVKR—DVYEEDSYVKR (3)(4)<br>DVYEEDSYVKR—DVYEEDSYVK (3)(4)(5)<br>ISDFGLSRDVYEEDSYVKR—DVYEEDSYVKR (3)(4)(5) | 18 | - | Yes | Yes |
| K96— K722 | KASVTIEDPK—ASVTIEDPKWEFPRK (5) | 1 | Yes | Yes | - |
| K71— K71 | LASLQQENKVLK—LASLQQENKVLK (3)(4)(5)<br>LASLQQENKVLK—LASLQQENK (4) | 5 | Yes | Yes | Yes |
| K722— K728 | KNLVLGK—NLVLGKTLGEGEFGK (3)(4) | 2 | - | - | - |
| K722— K737 | KNLVLGK—NLVLGKTLGEGEFGK (3) | 1 | - | Yes | - |
| K747— K747 | ATAFHLKGR— ATAFHLKGR (5)<br>ATAFHLKGR— TLGEGEFGKVVKATAFHLK (4) | 2 | Yes | - | - |
| S98— K716 | KASVTIEDPK—ASVTIEDPKWEFPR (4)<br>ASVTIEDPKWEFPR—KASVTIEDPK (4) | 2 | Yes | Yes | Yes |
| K740— K747 | TLGEGEFGKVVK—TLGEGEFGKVVKATAFHLK (5) | 1 | - | - | - |
| T100— K716 | KASVTIEDPK—ASVTIEDPKWEFPR (4) | 4 | Yes | | Yes |
| K722— K722 | KNLVLGK— KNLVLGK (3)(4)<br>ASVTIEDPKWEFPRK—KNLVLGK (5) | 4 | Yes | - | - |
| K1003— K1011 | RPVFADISKDLEK— DLEKMMVK (4) | 1 | - | - | - |
| K737— K737 | NLVLGKTLGEGEFGK— TLGEGEFGKVVK (4) | 1 | - | Yes | - |
| K728— K740 | NLVLGKTLGEGEFGK—TLGEGEFGKVVK (4) | 1 | - | - | - |
| K737— K740 | TLGEGEFGK—TLGEGEFGKVVK (3) | 2 | - | - | - |
| K1060— K1060 | ALPSTWIENKLYGR— ALPSTWIENKLYGR (3)(4)(5)<br>ALPSTWIENKLYGR— ALPSTWIENK (4) | 7 | - | - | - |
| K716— K716 | ASVTIEDPKWEFPR— ASVTIEDPKWEFPR (4) | 1 | Yes | - | Yes |
| S98— S98 | ASVTIEDPK— ASVTIEDPK (3) | 1 | - | Yes | Yes |
| K716— K722 | ASVTIEDPKWEFPRK—KNLVLGK (4) | 1 | Yes | - | Yes |
| K740— T742 | TLGEGEFGKVVK—VVKATAFHLK (5) | 1 | - | - | - |
| K71— K74 | LASLQQENKVLK—VLKIELETYK (4) | 1 | Yes | Yes | Yes |

Supplemental table 4. Intermolecular crosslinked peptides from the XL-MS analyses for CCDC6-RET apo state filtered with the 3D-structural models: model 1 (active like, *cis* autophosphorylation), model 2 (inactive, face-to-face transinhibited) and model 3 (transition state). Highlighted crosslinked peptides are found exclusively in the apo state, see figure 8.

| Crosslinked residues | Crosslinked peptides | Abundance in 0P state | Model #1 | Model #2 | Model #3 |
|---|---|---|---|---|---|
| K737— K737 | TLGEGEFGKVVK—TLGEGEFGKVVK (3)(4)<br>NLVLGKTLGEGEFGK— TLGEGEFGKVVK (4)<br>NLVLGKTLGEGEFGK— NLVLGKTLGEGEFGK (4) | 6 | - | Yes | - |
| K737— K747 | TLGEGEFGKVVK—TLGEGEFGKVVKATAFHLK (5) | 1 | - | - | - |
| S98— K716 | KASVTIEDPK—ASVTIEDPKWEFPR (4) | 3 | Yes | Yes | Yes |
| K71— K74 | LASLQQENKVLK—VLKIELETYK (4) | 1 | - | - | - |
| K747— K747 | ATAFHLKGR— ATAFHLKGR (4)(5)<br>VVKATAFHLK— VVKATAFHLK (5)(6)<br>NLVLGKTLGEGEFGKVVKATAFHLK— VVKATAFHLK (5)(7)<br>VVKATAFHLKGR— ATAFHLK (5) | 14 | Yes | - | - |
| K96— K96 | KASVTIEDPK— KASVTIEDPK (4) | 2 | Yes | - | Yes |
| K722— K737 | KNLVLGK—NLVLGKTLGEGEFGK (3)(4) | 2 | - | Yes | - |
| K728— K737 | NLVLGKTLGEGEFGK—TLGEGEFGKVVK (3)<br>KNLVLGKTLGEGEFGK—TLGEGEFGK (4) | 2 | - | - | - |
| K722— K728 | KNLVLGK—NLVLGKTLGEGEFGK (3)(4)(5) | 3 | - | - | - |
| K740— K747 | TLGEGEFGKVVK—VVKATAFHLK (5)<br>NLVLGKTLGEGEFGKVVKATAFHLK—VVKATAFHLK (5)(6)<br>TLGEGEFGKVVK—VVKATAFHLKGR (5)<br>VVKATAFHLK—ATAFHLKGR (5)<br>TLGEGEFGKVVK—TLGEGEFGKVVKATAFHLK (4)(5)<br>NLVLGKTLGEGEFGKVVKATAFHLK—VVKATAFHLK (5) | 12 | - | - | - |
| K740— T742 | TLGEGEFGKVVK—TLGEGEFGKVVKATAFHLK (5) | 1 | - | - | - |
| K737— K740 | TLGEGEFGKVVK—VVKATAFHLK (5)<br>TLGEGEFGKVVKATAFHLK—TLGEGEFGKVVK (4)<br>NLVLGKTLGEGEFGK—TLGEGEFGKVVK (4)<br>TLGEGEFGK—TLGEGEFGKVVK (3) | 5 | - | - | - |
| T100— K716 | KASVTIEDPK—ASVTIEDPKWEFPR (3)(4) | 3 | Yes | - | Yes |
| T742— T742 | ATAFHLKGR— ATAFHLKGR (3) | 1 | - | - | - |
| K722— T729 | KNLVLGKTLGEGEFGKVVK—TLGEGEFGK (4) | 2 | - | - | - |
| K96— K722 | KASVTIEDPK—ASVTIEDPKWEFPRK (5) | 1 | Yes | Yes | - |
| K740— K740 | VVKATAFHLK—VVKATAFHLK (5)(6)<br>TLGEGEFGKVVK—TLGEGEFGKVVK (5)<br>NLVLGKTLGEGEFGKVVKATAFHLK—VVKATAFHLK (5)(6)(7)<br>TLGEGEFGKVVKATAFHLK—TLGEGEFGKVVK (5)<br>TLGEGEFGKVVK—VVKATAFHLK (3)(4) | 14 | - | - | - |
| K1007— K1007 | DLEKMMVK— DLEKMMVK (4) | 1 | - | - | - |
| K1003— K1007 | RPVFADISKDLEK—DLEKMMVK (4) | 1 | - | - | - |
| K716— K722 | ASVTIEDPKWEFPRK—KNLVLGK (4)<br>ASVTIEDPKWEFPR—ASVTIEDPKWEFPRK (5) | 5 | Yes | - | Yes |
| K728— K740 | NLVLGKTLGEGEFGK—TLGEGEFGKVVK (4) | 1 | - | - | - |
| K989— K994 | LMoxLQCcmWKQEPDK—LMLQCcmWK (3)(4)<br>MERPDNCcmSEEMYRLMLQCcmWK—LMoxLQCcmWKQEPDK (5) | 5 | - | - | - |
| K821— K821 | KVGPGYPDER—KVGPGYPDER (3)(4) | 3 | - | - | - |
| K722— K722 | KNLVLGK—KNLVLGK (2)(3)(4)<br>ASVTIEDPKWEFPRK— KNLVLGK (3)(5)<br>ASVTIEDPKWEFPRK— ASVTIEDPKWEFPRK (5)(6) | 11 | Yes | - | - |
| K737— T742 | TLGEGEFGKVVK—VVKATAFHLK (4) | 1 | - | - | - |
| K716— K716 | ASVTIEDPKWEFPR— ASVTIEDPKWEFPR (3)(4)(5)<br>KASVTIEDPKWEFPR— ASVTIEDPKWEFPR (5) | 8 | Yes | - | Yes |
| K728— K728 | NLVLGKTLGEGEFGK— NLVLGKTLGEGEFGK (4) | 1 | - | - | - |
| K1060— K1060 | ALPSTWIENKLYGR—ALPSTWIENKLYGR (3)(4) | 3 | - | - | - |
| K71— K71 | LASLQQENKVLK— LASLQQENKVLK (3)(4)(5) | 5 | Yes | Yes | Yes |
| K96— K716 | KASVTIEDPK—ASVTIEDPKWEFPRK (4)(5) | 3 | Yes | Yes | Yes |

Supplemental table 4 (continuation). Intermolecular crosslinked peptides from the XL-MS analyses for CCDC6-RET phosphorylated state filtered with the 3D-structural models: model 1 (active like, cis autophosphorylation), model 2 (inactive, face-to-face transinhibited) and model 3 (transition state). Highlighted crosslinked peptides are found exclusively in the phosphorylated state, see figure 8.
